# Network communication models improve the behavioral and functional predictive utility of the human structural connectome

**DOI:** 10.1101/2020.04.21.053702

**Authors:** Caio Seguin, Ye Tian, Andrew Zalesky

## Abstract

The connectome provides a structural substrate facilitating communication between brain regions. We aimed to establish whether accounting for polysynaptic communication paths in structural connectomes would improve prediction of interindividual variation in behavior as well as increase structure-function coupling strength. Structural connectomes were mapped for 889 healthy adults participating in the Human Connectome Project. To account for polysynaptic signaling, connectomes were transformed into communication matrices for each of 15 different network communication models. Communication matrices were (i) used to perform predictions of five data-driven behavioral dimensions and (ii) correlated to interregional resting-state functional connectivity (FC). While FC was the most accurate predictor of behavior, network communication models, in particular communicability and navigation, improved the performance of structural connectomes. Accounting for polysynaptic communication also significantly strengthened structure-function coupling, with the navigation and shortest paths models leading to 35-65% increases in association strength with FC. Combining behavioral and functional results into a single ranking of communication models positioned navigation as the top model, suggesting that it may more faithfully recapitulate underlying neural signaling patterns. We conclude that network communication models augment the functional and behavioral predictive utility of the human structural connectome and contribute to narrowing the gap between brain structure and function.

## INTRODUCTION

Communication between gray matter regions is crucial for brain functioning. The complex topology of the structural connectome [1, 2] provides the scaffold on top of which neural signaling unfolds. While region pairs that share a connection in the structural connectome may communicate directly, polysynaptic paths comprising two or more connections are required to establish communication between anatomically unconnected regions. Network communication models describe strategies to delineate putative signaling paths between regions, based on aspects of connectome topology and geometry [3].

Several network communication models have been proposed to describe large-scale neural signaling, ranging from naive random walk processes to optimal routing via shortest paths [4]. By considering polysynaptic paths, these models quantify the putative efficiency of communication between both connected and unconnected nodes, thus enabling a high-order structural description of interactions among every pair of regions in the connectome [5]. Recent studies report that network communication models can improve the strength of coupling between structural and functional connectivity in the human connectome [6], explain established patterns of cortical lateralization [7], and infer the directionality of effective connectivity from structural connectomes [8].

Building on this previous work, we aimed to investigate the utility of a range of candidate models of network communication in structural brain networks. First, we sought to determine whether accounting for polysynaptic (multi-hop) paths in structural brain networks using models of network communication would: i) improve the prediction of interindividual variation in behavior, compared to predictions based on direct structural connections alone; and, ii) improve the strength of structure-function coupling. Second, we aimed to establish a ranking of communication models with respect to their predictive utility, with the goal of determining which models may more faithfully capture biological signaling patterns related to behavior and FC.

We considered five previously proposed communication measures: (i) shortest paths [9, 10], (ii) navigation [11, 12], (iii) diffusion [13], (iv) search information [6, 14], and (v) communicability [15-17]. Collectively, these models cover a wide-range of neural signaling conceptualizations. Shortest paths and navigation deterministically route information using centralized and decentralized strategies, respectively. In contrast, diffusion and search information model communication from the stochastic perspective of random walk processes. Finally, communicability implements a broadcasting model of signaling, in which signals are simultaneously propagated along multiple network fronts. While all these candidate models have been investigated in the human connectome, which particular models provide the most parsimonious representation of large-scale neural signaling remains unclear.

Using diffusion-weighted MRI and tractography, we mapped structural connectivity (SC) matrices for *n* = 889 healthy adults participating in the Human Connectome Project (HCP) [18]. Each individuals SC matrix was then transformed into a communication matrix, which represented the efficiency of communication between each pair of regions under a particular candidate model of network communication. For each model, communication matrices were fed to statistical techniques to perform out-of-sample prediction of individual variation in five behavioral dimensions [19], and also correlated with FC matrices mapped using resting-state functional MRI. This enabled a systematic ranking of network communication models in terms of behavior prediction and structure-function coupling. While these criteria do not constitute direct biological validation of signaling strategies, we hypothesize that the higher the predictive utility of a communication model, the more likely it is to parsimoniously recapitulate the signaling mechanisms of the human brain.

## RESULTS

### Brain network communication matrices

Structural connectomes were mapped using white matter tractography applied to diffusion MRI data acquired for 889 healthy adults participating in the Human Connectome Project [18] (*Materials and Methods*). We focus on reporting results for connectomes comprising *N* = 360 cortical regions [20] that were thresholded to eliminate potentially spurious connections [21]. Results for alternative cortical parcellations and connection density thresholds are reported in the *Supplementary Information.*

Connectome mapping yielded a structural connectivity (SC) matrix for each individual. These matrices represented connectivity between directly connected regions and were generally sparse due to an absence of white matter tracts between a majority of region pairs. To model the impact of polysynaptic neural signaling, each individual’s connectivity matrix was transformed into a communication matrix (Fig. 1a). Communication matrices were of the same dimension as the SC matrices, but fully connected in most cases, and they quantified the efficiency of communication between indirectly (polysynaptic) as well as directly connected pairs of regions under a given network communication model. In contrast, the SC matrices only characterized directly connected pairs of regions.

**FIG. 1.**
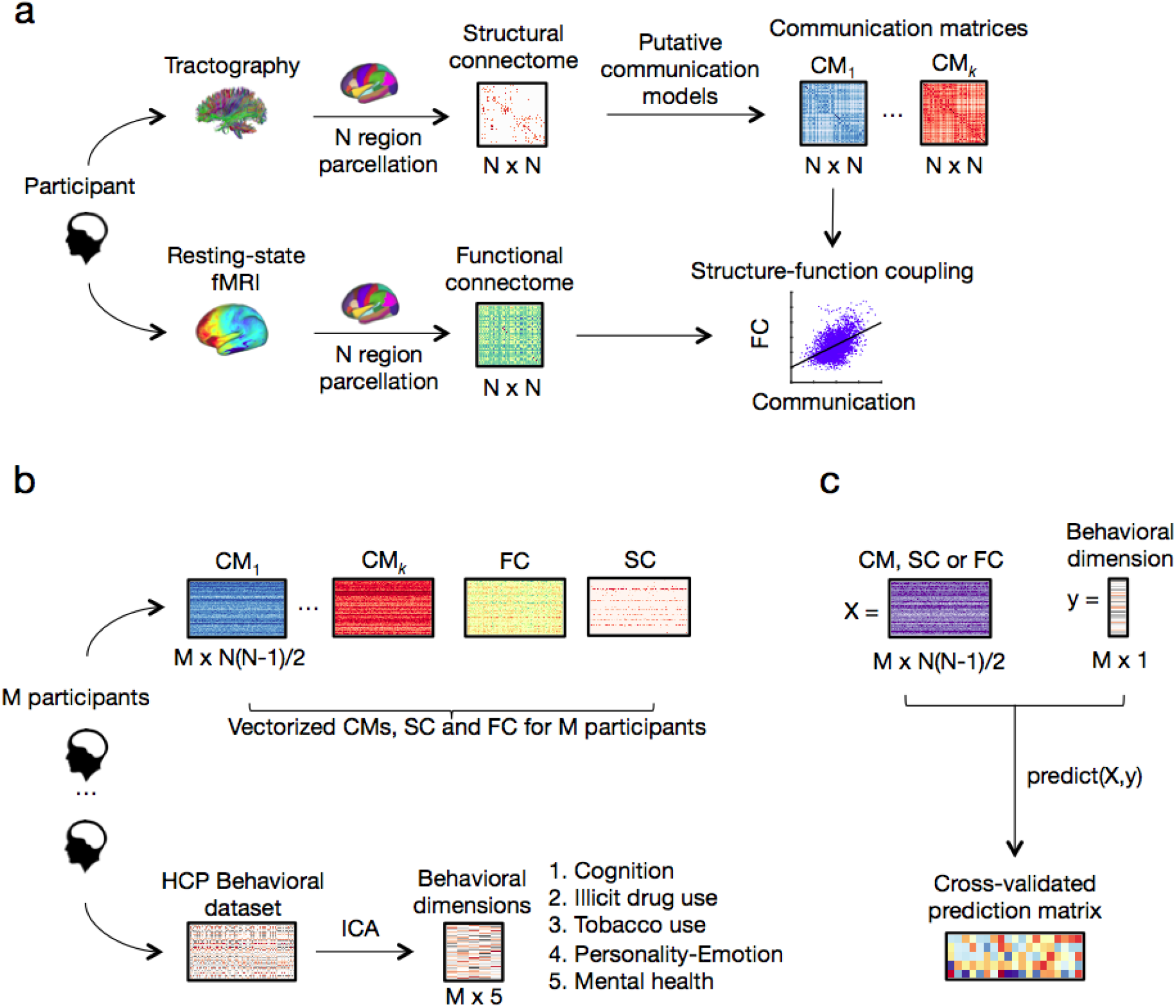
Methodology overview. **(a)** For each participant in our sample, structural connectomes comprising *N* cortical regions were mapped using white matter tractography applied to diffusion MRI. Structural connectivity matrices were transformed into communication matrices 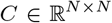, where *C*(*i, j*) denotes the communication efficiency from region *i* to region *j*. For each participant, a total of 15 communication matrices were derived representing different combinations of network communication models (shortest paths, navigation, diffusion, search information, communicability) and connection weight definitions (binary, weighted, distance). To assess structure-function coupling, communication matrices were correlated with FC matrices computed from resting-state functional MRI data. **(b)** Communication, FC and SC matrices were vectorized and aggregated across *n* = 889 participants, resulting in 17 *n* × *N*(*N* − 1)/2 matrices of explanatory variables. A set of five behavioral dimensions was computed by applying independent component analysis (ICA) to the HCP dataset of behavioral phenotypes. **(c)** Communication, SC and FC matrices were used to predict behavior. An entry in the resulting 17 × 5 prediction matrix corresponds to the mean cross-validated association between a communication or connectivity matrix and a behavioral dimension.

We considered three connectivity weight definitions: (i) *Weighted:* connection weights defined as the number of tractography streamline counts between regions; (ii) *Binary:* non-zero connection weights set to one; and (iii) *Distance:* non-zero connection weights set to the Euclidean distance between regions. Network communication models computed on these connectomes operationalize metabolic factors conjectured to shape large-scale signaling: (i) adoption of high-caliber, high-integrity white matter projections to enable fast and reliable signal propagation (weighted); (ii) reduction of the number of synaptic crossings (binary); and (iii) reduction of the physical length traversed by signals (distance).

### Predicting behavior with models of connectome communication

Statistical models were trained to independently predict five dimensions of behavior (cognition, illicit substance use, tobacco use, personality-emotional traits, mental health) based on features comprising an individual’s communication matrix (Fig. 1b,c). Training and prediction were performed separately for a total of 15 communication matrices representing different connection weight definitions (binary, weighted, distance) and network communication models (shortest paths, navigation, diffusion, search information, communicability). Additionally, predictions based on an individual’s SC and FC were computed to provide a benchmark. The five behavioral components represent orthogonal dimensions that were parsed from a comprehensive set of behavioral measures using independent component analysis (*Materials and Methods*).

Out-of-sample prediction accuracy was evaluated for 10 repetitions of a 10-fold cross-validation scheme. The Pearson correlation coefficient between the actual and out-of-sample predicted behavior was used to quantify prediction accuracy for each behavioral dimension. To ensure that our results were not contingent on the adoption of a particular statistical model, predictions were independently performed using lasso regression [22] and a regression model based on features identified by the network-based statistic (NBS) [23] (*Materials and Methods*). Prediction accuracies were averaged across cross-validation folds and repetitions, and visualized in the form of a matrix comprising behavioral dimensions (rows) and communication models (columns) (Fig. 2a,c).

**FIG. 2.**
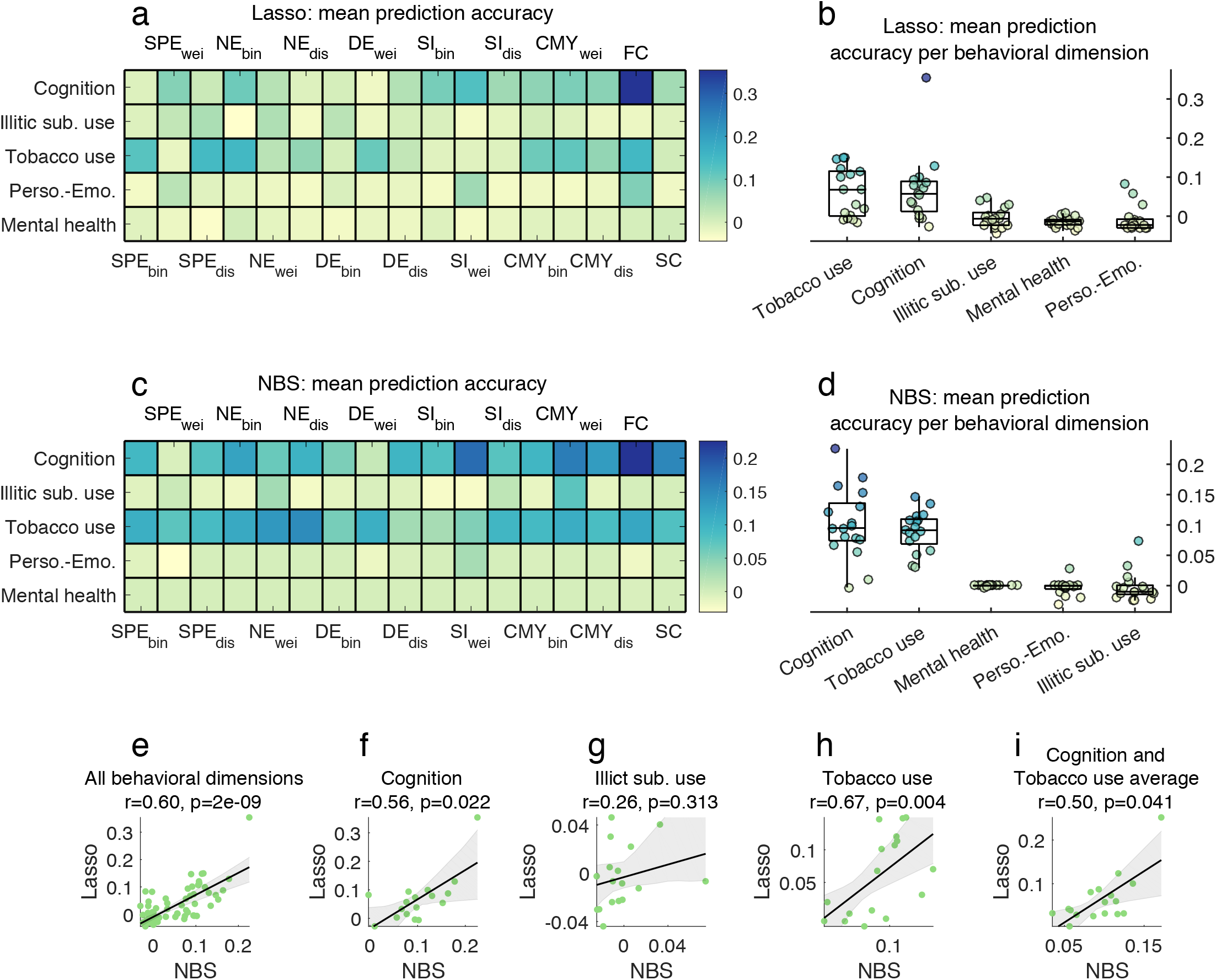
Predicting individual variation in human behavioral dimensions using models of connectome communication, as well as structural and functional connectivity (*N* = 360 thresholded connectomes). **(a)** Matrix of Pearson correlation coefficients between actual and out-of-sample predicted behavior using lasso regression for five orthogonal behavioral dimensions (rows) and 15 connectome communication models as well as SC and FC (columns, predictors). **(b)** Lasso regression prediction accuracies (correlation coefficients) stratified by behavioral dimensions. Each boxplot summarizes a row of the prediction accuracy matrix and the superimposed data points are colored accordingly. Top and bottom edges boxplots indicate, respectively, the 25th and 75th percentiles, while the central mark shows the distribution median. Mean prediction accuracies significantly differed between the five behavioral dimensions (*F*_(4,80)_ = 10.67, *p* = 5 × 10^−7^). **(c-d)** Same as (a-b), but for predictions carried out using a regression model based on featured identified by the NBS. Again, mean prediction accuracies were significantly different between behavioral dimensions (*F*_(4,80)_ = 47.18, *p* = 2 × 10^−20^). Scatter plots showing the correspondence (Spearman rank correlation coefficient and *p*-value) between lasso and NBS prediction accuracies for **(e)** all behavioral dimensions, **(f)** cognition, **(g)** illicit substance use, **(h)** tobacco use, and **(i)** the average between cognition and tobacco use prediction accuracies. SPE: shortest path efficiency, NE: navigation efficiency, DE: diffusion efficiency, SI: search information, CMY: communicability, bin: binary, wei: weighted, dis: distance.

We found that individual variation in some behavioral dimensions could be predicted with greater accuracy than others (lasso: *F*_(4,80)_ = 10.67, *p* = 5 × 10^−7^; NBS: *F*_(4,80)_ = 47.18, *p* = 2 × 10^−20^). Dimensions characterizing cognition (respective lasso and NBS accuracies averaged across all predictors: 0.068, 0.101) and tobacco use (0.061, 0.089) could be predicted more accurately on average, whereas comparably weaker predictions of illicit substance use (−0.003, −0.002), personality-emotion (−0.008, −0.003) and mental health (−0.014, −0.0003) were evident (Fig. 2b,d).

Prediction accuracies were consistent between the two statistical models (NBS, lasso), both when pooling the five behavioral dimensions (Spearman rank correlation coefficient *r*_(83)_ = 0.60, *p* = 2 × 10^−9^; Fig. 2e), as well as separately for cognition (*r*_(16)_ = 0.56, *p* = 0.022; Fig. 2f) and tobacco use (*r*_(16)_ = 0.67, *p* = 0.004; Fig. 2h). However, lasso and NBS diverged for the dimensions that were less accurately predicted (e.g., *p* = 0.313; Fig. 2g).

Focusing on lasso regression, we sought to determine whether behavioral predictions were robust to variations in our methodological settings. First, we found that adopting the mean square error to quantify predictive utility led to accuracies significantly associated to the ones computed based on Pearson correlation (Fig. S1). Second, we tested whether prediction accuracies were sensitive to changes in our connectome mapping pipeline. To this end, we recomputed behavioral predictions for three additional sets of connectomes: (i) *N* = 360 regions without connection thresholding, (ii) *N* = 68 regions with connection thresholding, and (iii) *N* = 68 regions without connection thresholding (*Materials and Methods*). Prediction accuracies were typically significantly correlated across low- and high-resolution, as well as thresholded and unthresholded, connectomes (Fig. S2). This was the case both when considering all behaviors and when restricting the analyses to the cognition and tobacco use dimensions.

Together, these findings suggest that network communication models (as well as SC and FC) can explain out-of-sample inter-individual variance in behavior. More specifically, cognition and tobacco use were the most accurately and robustly predicted behavioral dimensions. For this reason, we henceforth focus subsequent analyses on the averaged prediction accuracy obtained for the cognition and tobacco use dimensions. This provides us with a single measure of how connectome communication relates to behavior by considering only the behavioral traits that can be predicted with relevant accuracy. The obtained prediction accuracy average was also consistent across the lasso and NBS (*r*_(16)_ = 0.50, *p* = 0.041; Fig. 2i).

### Communication models improve the behavioral predictive utility of the human connectome

We sought to compare communication models, as well as SC and FC, in terms of their behavioral prediction accuracy. Figure 3a shows the distributions of out-of-sample accuracies (10 repetitions of 10-fold cross validation, averaged for the cognition and tobacco use dimensions) obtained for the each predictor using lasso regression. Accuracy distributions were ranked based on their medians. FC (median accuracy: 0.24) provided markedly greater prediction accuracy than all communication models and SC. Binary navigation (median accuracy: 0.12) and weighted communicability (median accuracy: 0.10) followed as the second and third most predictive communication models. Crucially, we observed that majority of communication models yielded greater prediction accuracy than SC (median accuracy: 0.03). This indicates that modeling polysynaptic signaling through the transformation of SC into communication matrices improved the behavioral predictive utility of structural connectomes.

**FIG. 3.**
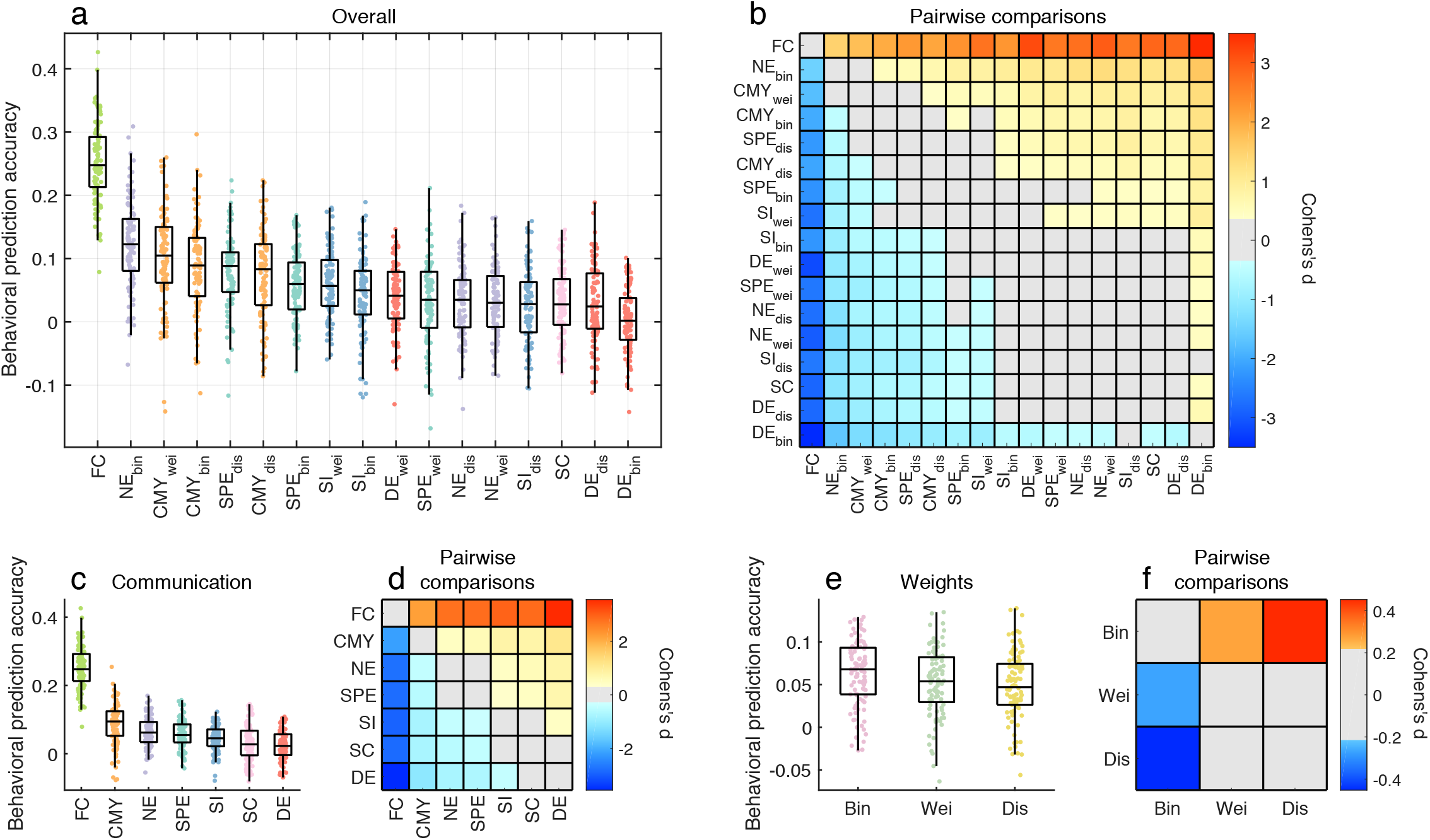
Comparison of the behavioral predictive utility of connectome communication models (Lasso regression, *N* = 360 thresholded connectomes, average cognition and tobacco use prediction accuracies). Across panels, top and bottom edges boxplots indicate, respectively, the 25th and 75th percentiles, while the central mark shows the distribution median. **(a)** Prediction accuracy distributions for 10 repetitions of 10-fold cross-validation. Communication models, SC and FC were sorted based on their median prediction accuracy. **(b)** Effect size matrix of pairwise statistical comparisons between predictors. Warm- and cool-colored cells indicate predictor pairs with significantly different means, as assessed by a repeated-measures t-test (Bonferroni-corrected for 136 multiple comparisons with significance threshold *α* = 3.67 × 10^−4^). A warm-colored *i, j* matrix entry indicates that predictor *i* yields significantly more accurate predictions than predictor *j*. **(c)** Prediction accuracy distributions of communication models averaged across connection weight definitions. SC and FC were not subjected to any averaging and accuracies remain the same as in panel (a). **(d)** Effect size matrix of pairwise repeated-measures t-tests between distributions in panel (c), with colored cells indicating significant differences in mean prediction accuracies (Bonferroni-corrected with significance threshold *α* = 0.0024). **(e)** Prediction accuracy distributions of connectomes with different connection weight definitions averaged across communication models. **(f)** Same as panel (d), but Bonferroni-corrected with significance threshold *α* = 0.0167.

We performed repeated measures t-tests to assess pairwise statistical differences in the predictive utility of communication models and connectivity measures (*Materials and Methods*). Figure 3b shows the effect size matrix (Cohen’s d; Bonferroni-corrected for 136 multiple comparisons with significance threshold *α* = 3.67 × 10^−4^) of differences between mean prediction accuracies, with warm- and cool-colored cells indicating model pairs for which a significant difference was observed. As expected, FC outperformed all other predictors (e.g., *p* = 1 × 10^−26^ between FC and binary navigation), while there was no evidence for a difference in predictive utility between binary navigation (2nd best predictor) and weighted communicability (3rd best predictor) following correction for multiple comparisons (*p* = 0.002). The lack of colored cells along the main diagonal of the effective size matrix indicates that predictors of similar ranking seldom yielded significantly different accuracy. Importantly, seven communication models (out of 15) significantly outperformed SC, including binary navigation; binary, weighted and distance communicability; binary and distance shortest paths; and weighted search information (all *p* < 10^−4^). This underscores the improvement in behavioral predictive utility gained from accounting for polysynaptic communication in structural connectomes, compared to predictions that only account for direct structural connections.

Next, we aimed to separate the effects of communication model choice and connection weight definition on prediction accuracy. To this end, prediction accuracies were averaged over the three weight definitions for each communication model (Fig. 3c,d), or averaged over the 15 models for each weight definition (Fig. 3e,f). Prediction accuracies for FC and SC remained the same. With respect to the effect of communication model, we found that communicability significantly outperformed other communication models and SC (e.g., *p* = 3 × 10^−5^, 2 × 10^−11^ for comparisons of communicability to navigation and SC, respectively), although FC remained the leading predictor. Navigation and shortest paths featured in second and third positions, both performing better than SC (*p* = 3 × 10^−7^, 3 × 10^−5^, respectively) and with no statistical difference between them (*p* = 0.26).With respect to connection weight definition binary connectomes yielded significantly higher prediction accuracies, on average, compared to weighted and distance connectomes (*p* = 0.009, 2 × 10^−5^, respectively), albeit with a weaker effect size than differences between communication models. This suggests that the choice of communication model may be more important to behavior predictions than the definition of connection weights.

In order to assess the robustness of these results, we executed additional analyses in which we (i) substituted the lasso prediction model with a regression model applied to features identified by the NBS (Fig. S3) and (ii) considered predictions for the cognition and tobacco use dimensions separately (*Supplementary Note 1*; Figs. S4, S5, S6, S7). The NBS prediction method largely reticulated lasso results (as was indicated by Fig. 2e-i). FC remained the strongest predictor of behavior, although with a smaller margin of difference to navigation and communicability communication models. Examining cognition and tobacco use independently reiterated the overall good performance of navigation and communicability. Importantly, however, it also revealed the presence of certain dimension-specific relationships between communication and behavior. For instance, search information yielded top- and bottom-ranking predictions for cognition and tobacco use, respectively.

Taken together, the behavioral prediction analyses led to three key findings. First, behavioral predictions were more accurate when performed based on functional rather than structural attributes. Second, while navigation and communicability typically showed high predictive utility, our results did not point towards a single communication model as the best predictor of human behavior. This indicates that different communication models may be better suited to predict different behavioral dimensions, possibly suggesting the presence of behavior-specific signaling mechanisms in the human brain. Third, and most importantly, the transformation of SC (only direct connections) into communication matrices (models of polysynaptic interactions) led to an improvement of structural-based predictions, bringing them closer to the predictive utility of FC. Collectively, these findings indicate that connectome communication models capture higher-order structural relations among brain regions that can better account for interindividual variation in behavior than SC alone.

### Communication models improve structure-function coupling

We next investigated whether accounting for network communication in the structural connectome can improve the strength of the relation between SC and FC, known as structure-function coupling. Classically, associations have been directly tested between structural and functional connections [24]. A growing body of work indicates that accounting for higher-order regional interactions through models of polysynaptic signaling (i.e., transforming structural connectomes into communication matrices) can improve structure-function coupling [5, 6, 8, 25, 26]. For two regions that are not directly connected with an anatomical fiber, strong FC is conjectured to indicate the presence of an efficient signaling path that facilitates communication through the underlying anatomical connections [3].

To test this hypothesis, we computed the association between FC and communication matrices for each individual in our sample. Additionally, as a benchmark, we also considered the association of FC to SC and to interregional Euclidean distance. Associations were computed as the Spearman correlation between upper triangular matrix entries. In addition to individual-level associations, we also analyzed structure-function coupling derived from group-level SC and FC. Finally, associations were derived for coarse- (*N* = 68 regions) and fine-grained (*N* = 360 regions) connectomes, which were thresholded prior to the computation of communication models. FC matrices were not thresholded. Further details on the computation of structure-function coupling are provided in the *Materials and Methods*.

As previously reported [6], communication matrices were correlated with FC, irrespective of the particular communication model (Fig. 4). In other words, FC was generally stronger between regional pairs interconnected by more efficient communication pathways. Group-level correlations (*r_G_*; black crosses) were universally stronger than those obtained for the median individual (**r*_I_*; box-plots), supporting the notion that predicting population-level FC traits is less challenging than modeling idiosyncratic relationships between brain structure and function.

**FIG. 4.**
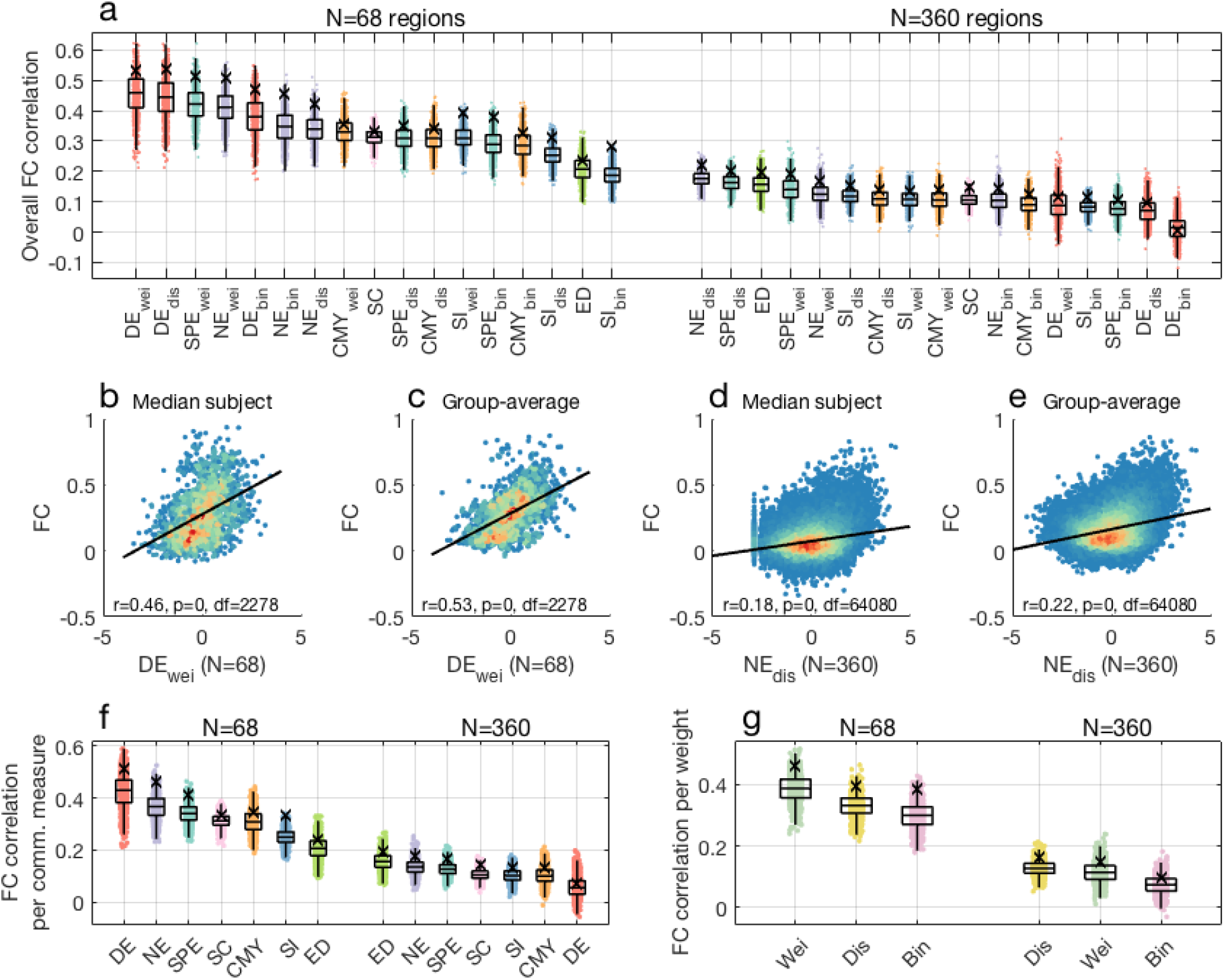
Structure-function coupling across connectome communication models (*N* = 68, 360 thresholded connectomes). **(a)** Data points show individual-level correlation of FC to communication, SC and Euclidean distance matrices. Black crosses indicate correlations obtained for group-averaged matrices. Top and bottom edges boxplots indicate, respectively, the 25th and 75th percentiles, while the central mark shows the distribution median. Communication models, SC and Euclidean distance were ranked according to the median structure-function coupling strength across individuals. **(b)** Scatter plot depicting the relationship between FC and the top-ranked communication model for connectomes comprising *N* = 68 regions, for the median individual. For ease of visualization, communication matrix entries were resampled to normal distributions. Warm and cool colors indicate high and low data point density, respectively. **(c)** Same as (b), but for group-average matrices. **(d-e)** Same as (b-c), but for connectomes comprising *N* = 360 regions. **(f)** Structure-function coupling for communication models, SC and Euclidean distance, averaged across connection weight definitions. **(g)** Structure-function coupling obtained for binary, weighted and distance connectomes, averaged across communication models.

**FIG. 5.**
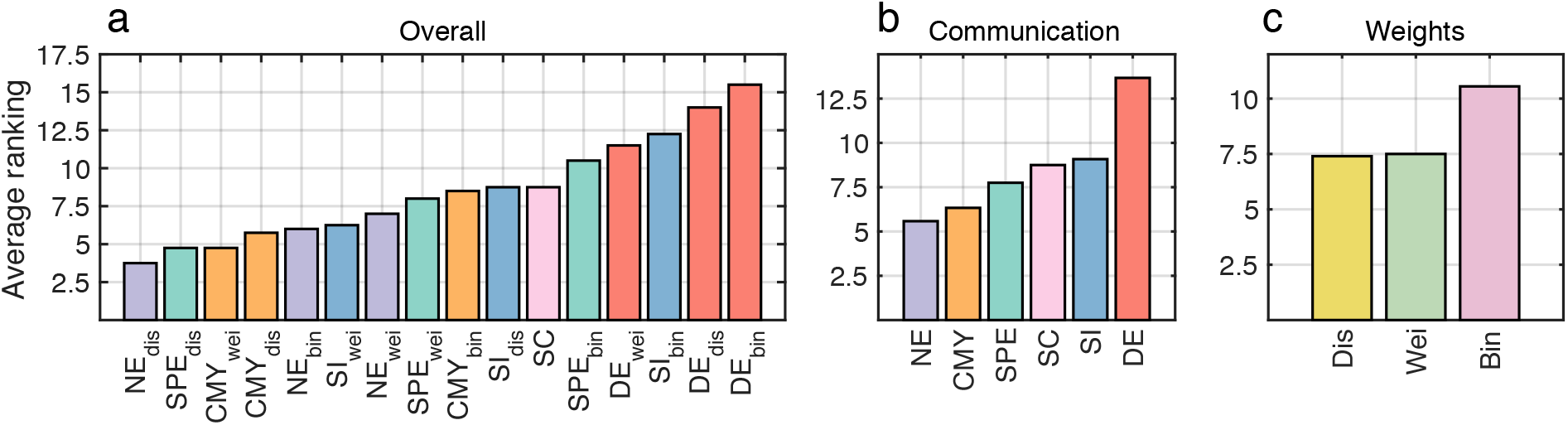
Overall ranking of communication models (*N* = 360 thresholded connectomes). Communication models and SC were ranked according to averaged behavioral and functional predictions. **(a)** Overall predictive utility ranking for *N* = 360 thresholded connectomes. Ranking shown in (a) grouped by **(b)** communication models and **(c)** connection weight definitions.

We found that parcellation resolution had a strong influence on the strength of structure-function coupling. The link between structure and function weakened for high resolution connectomes, irrespective of the communication model (Fig. 4a). Moreover, the ranking of communication models in terms of structure-function coupling differed between connectome resolutions (Spearman rank correlation between low- and high-resolution FC predictions *p* = 0.65). For *N* = 68 regions, weighted and distance diffusion yielded the strongest structure-function couplings (**r*_I_* = 0.46 and **r*_G_* = 0.53 for weighted diffusion; Fig. 4b,c). This recapitulates previous work indicating the functional predictive utility of random walk models applied to connectomes comprising less than one hundred regions [25]. However, in sharp contrast, diffusion performed poorly for *N* = 360 regions, going from yielding the most accurate estimates of FC in low-resolution but ranking as the worst overall predictor in high-resolution. Conversely, the coupling between Euclidean distance and FC showed the opposite relationship to connectome resolution, with interregional distances leading to weak and strong associations for coarse- and fine-grained parcellations, respectively.

Navigation and shortest paths resulted in consistently high-ranked FC predictions regardless of connectome resolution. For *N* = 68 regions, weighted navigation and shortest paths showed comparable associations to the top-ranking diffusion models (e.g., *r_I_* = 0.42 for weighted shortest paths). For *N* = 360 regions, distance navigation was the top-ranking model (**r*_I_* = 0.18 and **r*_G_* = 0.22; Fig. 4d,e), followed by distance shortest paths in second place, both outperforming the Euclidean distance benchmark in the third position.

Crucially, despite the effects of connectome resolution, modeling polysynaptic communication on top of structural connectomes tightened structure-function coupling. This was the case for 8 and 9 out of the 15 communication models considered, for low- and high-resolution connectomes, respectively. For instance, for the median individual, weighted diffusion in 68-region connectomes strengthened coupling by 46% compared to SC, while computing distance navigation in 360-region connectomes boosted FC predictions by 66% compared to SC.

Grouping couplings by communication models reiterated functional predictive utility differences between low- and high-resolution connectomes (Fig. 4f). Grouping couplings by connection weight definitions showed that, on average, communication models computed on weighted and distance connectomes led to stronger couplings for coarse- and fine-grained parcellations, respectively (Fig. 4g), suggesting that the established influence of interregional distance in SC and FC [27, 28] may be stronger for connectomes derived at finer levels of areal granularity.

In summary, we observed that structure-function coupling is affected by connectome resolution and by whether associations are computed on individual or population levels. Regardless of parcellation granularity, most connectome communication models contributed to strengthening structure-function coupling. Moreover, navigation and shortest paths yielded the most accurate and reliable predictions of FC. While here we focused on thresholded connectomes, similar results were observed for unthresholded networks (Fig. S8). Rankings of functional predictive utility also remained consistent when stratifying analyses between structurally connected and unconnected region pairs, as well as for intrahemispheric structure-function associations (*Supplementary Note 2*; Fig. S9). Together, these observations build on the behavioral prediction findings, further supporting the notion that connectome communication models contribute to bridging the gap between brain structure and function.

### Ranking communication models

Finally, we aimed to derive a single ranking of predictive utility, as the average of behavioral and functional prediction accuracy rankings, for the 15 connectome communication models explored and SC (*Materials and Methods*). First, we focused on results obtained for thresholded connectomes comprising *N* = 360 regions. To this end, we averaged the rankings from three analyses: lasso behavioral predictions (Fig. 3), NBS behavioral predictions (Fig. S3) and FC predictions (Fig. 4a). We found that distance navigation showed the highest combined behavioral and functional predictive utility (average ranking *τ* = 3.7; 5a), followed by a tie between distance shortest paths and weighted communicability (*τ* = 4.7). SC featured in the 11th position and was outranked by most navigation, communicability, shortest paths and search information models. We also found that navigation was the top ranking model regardless of connection weight definitions (*τ* = 5.9; 5b), followed by communicability (*τ* = 6.3), shortest paths (*τ* = 7.7), SC (*τ* = 8.7), search information (*τ* = 9.1), and diffusion in the last position (*τ* = 13.6). Moreover, connection weights defined in terms of streamline counts and Euclidean distance resulted in similar predictive utility rankings (*τ* = 7.4, 7.5, respectively) while binary connectomes typically led to less accurate predictions (*τ* = 10.5).

Next, we computed rankings for the three alternative sets of connectomes considered in this study, namely *N* = 360 unthresholded, *N* = 68 thresholded and *N* = 68 unthresholded connectomes (Fig. S10). Contrasting results across these scenarios revealed that sparse and fine-grained connectomes benefited more from models of polysynaptic communication than dense and coarsegrained connectomes. Intuitively, a densely connected network requires few polysynaptic paths to propagate information, since most regions can communicate via direct connections. Consequently, while navigation, communicability and shortest paths typically improved the predictive utility of SC for *N* = 360, a markedly reduced number of signaling strategies was beneficial to the performance of connectomes comprising *N* = 68 regions. Importantly, across all sets of SC reconstructions, navigation was the only model that constantly featured as advantageous to the predictive utility of the human connectome.

## DISCUSSION

Human cognition and behavior arise from the orchestrated activity of multiple brain regions [29, 30]. Resisting-state FC is currently one of the most widely used neuroimaging measures to quantify this concerted activity [31–33]. It is thus unsurprising that statistical methods trained on functional brain networks led to the most accurate predictions of human behavior. Importantly, the signaling processes that facilitate synchronous interregional activity must unfold along structural connections forming direct or indirect (polysynaptic) communication paths. Therefore, brain structure, brain function, neural communication, and human behavior are tightly intertwined. This is corroborated by the key conclusion of the present study: accounting for polysynaptic communication in SC matrices can substantially improve structure-function coupling and the predictive utility of SC. Transforming a SC matrix into a communication matrix can be achieved efficiently for most communication models without the need for any sophisticated computational processing. We recommend performing the transformation when evaluating structure-function coupling or using SC to predict interindividual variation in behavior. While accounting for communication did not lead to SC outperforming FC with respect to behavior prediction, it narrowed that gap between the predictive utility of SC and FC.

As investigators tackle the longstanding challenge of elucidating the relationship between brain structure and function [34–36], it has become increasingly clear that FC arises from high-order regional interactions that cannot be explained by direct anatomical connections [5]. In line with this notion, we found that taking polysynaptic signaling into account through network communication models strengthened structure-function coupling. This observation recapitulates earlier reports on the functional predictive utility of connectome communication models [6] and provides support to the notion that FC is facilitated by communication pathways in the underlying structural connectome. Taken together, the behavioral and functional prediction analyses contribute empirical evidence that connectome communication models act as a bridge between structural and functional conceptualizations of brain networks [3, 37].

Importantly, brain structure-function relationships encompass a rich and diverse field of research, with several alternative classes of higher-order models showing promise in modeling function from structure. Examples include biophysical models of neural activity [38–40], statistical methods [41, 42], and other approaches centered around network communication that we did not explore in the present work [26, 43–45]. Likewise, relating neuroimaging data to behavior is a central goal of neuroscience [46, 47]. Recent studies have explored neural correlates of behavior and cognition by leveraging graph measures of brain organization [48, 49], dynamic patterns of FC fluctuations [50, 51], multivariate correlation methods [52, 53] and machine learning techniques [54, 55]. Our analyses sought to complement these efforts from the perspective of connectome communication. The goal of this paper was not to show that network communication models lead to more accurate predictions than alternative approaches, nor that our prediction scheme and statistical methods are superior to previously adopted techniques. Rather, we were interested in comparing the predictive utility of candidate models of connectome communication, as well as connectivity and distance benchmarks, in a controlled and internally consistent manner.

### Comparisons between connectome communication models

Communication matrices computed with the navigation and communicability models typically led to the highest-ranking behavioral predictions amongst the candidate signaling strategies explored. It is important to notice, however, that search information and shortest paths also performed well in certain scenarios. Therefore, while our behavioral results strongly suggested the benefits of modeling polysynaptic signaling, they did not provide a clear answer to the question of which communication models are most associated to human behavioral dimensions. Alternatively, our findings may indicate the interesting possibility that large-scale information integration in the brain is not facilitated by a unique signaling mechanism, and that different communication models may find more utility in describing varied behavioral and cognitive processes.

Navigation and shortest paths led to the most reliable FC predictions, featuring as the best models for high-resolution connectomes and closely following behind by diffusion for low-resolution connectomes. Navigation and shortest paths computed on distance connectomes led to FC predictions that surpassed those obtained from Euclidean distance, which exerts a well-documented influence on both SC and FC [27, 28, 56]. This indicates that combining the contributions of brain network topology and geometry might be helpful in modeling structure-function relationships. Furthermore, given the high efficiency of communication along navigation and shortest paths, these findings suggest that FC is facilitated primarily by highly efficient signaling pathways. This observation stands in contrast with previous work on the functional relevance of models that incorporate deviations from optimal routes, such as search information [6, 57]. We note, however, that navigation implements a decentralized identification of near-optimal paths [58, 59] that contributes to the understanding of how efficient signaling can be achieved in the absence of global knowledge of network architecture [12, 60].

Combining communication model rankings from behavioral and functional predictions revealed that navigation was the model with the highest overall predictive utility ranking. This finding provides initial evidence that, out of the putative communication strategies explored, navigation may most faithfully describe underlying neural signaling patterns supporting human behavior and brain function. Our results contribute to the growing body of work supporting the neuroscientific utility of network navigation [8, 12, 61–63].

In addition to investigating putative neural signaling strategies, we also considered different connection weight definitions. Polysynaptic transmission of neural signals entails metabolic expenditures related to the propagation of action potential along axonal projections and the crossing of synaptic junctions. Communication in the brain is thought to be metabolically frugal [30, 66], but what aspects of structural connectivity are relevant to energy consumption in large-scale signaling remain unclear. We found that weighted and distance connectomes typically led to communication matrices with higher predictive utility. This is initial evidence that neural signaling may favor communication paths prioritizing the adoption of physically short and high caliber connections, instead of paths that reduce the number of synaptic crossings between regions. Additionally, these observations warrant further investigation of the relatively unexplored distance connectome [67].

In accordance to previous reports [68, 69], we observed that FC predictions were more accurate for low-rather than high-resolution connectomes, as well as for group-rather than individual-level analyses. This is not surprising since the number of functional connections grows quadratically with the number of regions and capturing idiosyncrasies in FC is more challenging than modeling general principles of connectivity. Despite their simplicity, these observations are important to the validation of FC prediction methods, suggesting that models constructed and evaluated on coarse and population-level networks may not generalize to more challenging settings.

### Limitations and future directions

Several methodological limitations of the present work should be discussed. First, we reiterate that the main focus of our investigation was to perform an evaluation of candidate models of connectome communication. Although we explored multiple brain network reconstruction pipelines, we were not primarily concerned with which mapping techniques produced connectomes with the highest predictive utility. The choice of parcellation schemes [70] and whether or not to threshold structural connectomes [71, 72] are both complex open questions that fall outside the scope of this work. We also note that white matter tractography algorithms are susceptible to a number of known biases that could potentially impact our results [73].

Further research is necessary to untangle the contributions of specific brain regions to the predictive utility of different communication models. This could be achieved by examining lasso regression weights and NBS connected components. Alternatively, behavior and functional predictions could be performed based on regionwise communication efficiencies, rather than complete communication matrices [74]. Efforts in these directions could help elucidate how different communication models utilize features of connectome topology to facilitate information transfer.

While we sought to evaluate a wide-range of communication models, alternative network propagation strategies could provide valuable insight into mechanisms of neural signaling and warrant further research. These include linear transmission models [26], biased random walks [4], cooperative learning [75], dynamic communication models [76], and information-theoretic approaches [77].

In conclusion, we demonstrated that taking into account polysynaptic signaling via models of network communication improves the behavioral and functional predictive utility of the human structural connectome. Overall, navigation ranked as the communication model with highest predictive utility, indicating that it may faithfully approximate underlying signaling pathways related to human behavior and brain function. These findings contribute novel insights to researchers interested in characterizing information processing in nervous systems.

## ACKNOWLEDGMENTS

We thank Olaf Sporns for valuable discussions. Human data were provided by the Human Connectome Project, WUMinn Consortium (1U54MH091657; Principal Investigators: David Van Essen and Kamil Ugurbil) funded by the 16 National Institutes of Health (NIH) institutes and centers that support the NIH Blueprint for Neuroscience Research; and by the McDonnell Center for Systems Neuroscience at Washington University.

## MATERIALS AND METHODS

### Structural connectivity data

Minimally preprocessed high-resolution diffusion weighted magnetic resonance imaging (MRI) data was obtained from the Human Connectome Project (HCP) [18]. Details about the acquisition and preprocessing of diffusion MRI data are found in [78, 79]. Analyses were restricted to participants with complete HCP 3T imaging protocol, yielding a total sample of 889 healthy adults (age 22–35, 52.8% females). Whole-brain structural connectomes were mapped using diffusion tensor imaging and deterministic white matter tractography pipeline implemented using MRtrix3 [80] (FACT tracking algorithm, 5 × 10^6^ streamlines, 0.5 mm tracking step-size, 400 mm maximum streamline length and 0.1 fractional anisotropy cutoff for termination of tracks). The connection weight between a pair of regions was defined as the total number of streamlines connecting them, resulting in a *N* × *N* weighted connectivity matrix for each participant. Group-level structural connectomes were computed by averaging the connectivity matrices of all subjects.

We used cortical parcellations containing *N* = 68, 360 regions. The 68-region parcellation consists of the anatomically delineated cortical areas of the Desikan-Killiany atlas [81]. The 360-region parcellation is a multimodal atlas constructed from high-resolution structural and functional data from HCP [20]. We also considered thresholded and unthresholded connectomes. Following connection density thresholding, only the top 15% and 20% strongest connection (in terms of streamline counts) were kept in connectomes comprising 360 and 68 regions, respectively. A more lenient threshold was applied to connectomes with 68 regions in order to avoid network fragmentation. Unthresholded connectomes maintained all connections identified in the structural connectivity reconstruction process.

### Connection weight and length definitions

Structural connectomes can be defined in terms of *N* × *N* adjacency matrices of connectivity weights (*W*) or lengths (*L*). Connection weights provide a measure of the strength and reliability of anatomical connections between regions pairs, while connection lengths quantify the distance or travel cost between regions pairs. Different network communication measures are computed based on *W* (e.g., diffusion efficiency and communicability), *L* (e.g., shortest path efficiency and navigation efficiency) or a combination of both (e.g., search information).

We considered three definitions of *W*: weighted, binary and distance. In the weighted case, *W_wei_*(*i, j*) was defined as the total number of streamlines with one endpoint in region *i* and the other in region *j*. The binary adjacency matrix was defined as *W_bin_*(*i, j*) = 1 if *W_wei_*(*i, j*) > 0 and *W_bin_*(*i, j*) = 0 otherwise. Distance-based connectivity was defined as *W_dis_*(*i, j*) = 1/*D*(*i, j*) if *W_wei_*(*i, j*) > 0 and *W_dis_*(*i, j*) = 0 otherwise, where *D* is the Euclidean distance matrix between region centroids.

Similarly, *L* was also defined in terms of binary, weighted and distance connection traversal costs. In all three cases, *L*(*i, j*) = ∞ for *ij* region pairs that do not share a direct anatomical connection, ensuring that communication is restrict to unfold through the connectome. Binary (*L_bin_*) and distance-based (*L_dis_*) connection lengths are straightforwardly defined from their weight counterparts as *L_bin_*(*i, j*) = 1 if *W_bin_*(*i, j*) = 1 and *L_bin_*(*i, j*) = ∞ otherwise, and *L_dis_*(*i, j*) = *D*(*i, j*) if *W_dis_*(*i, j*) > 0 and *L_dis_*(*i, j*) = ∞ otherwise. Lengths based on the streamline count between regions pairs are defined by monotonic weight-to-length transformations that remap large connection weights into short connection lengths. This way, white matter tracts conjectured to have high caliber and integrity are considered faster channels of communication than weak and unreliable ones. We considered a logarithmic weight-to-length remapping such that *L_wei_* = −*log*_10_(*W_wei_*/*max*(*W_wei_*) + 1) (the unity addition to the denominator avoids the remapping of the maximum weight into zero length) [12], producing normally distributed lengths that attenuate the important of extreme weights [82, 83].

### Network communication models

In this section, we provide details regarding the definition of the five network communication models evaluated in this study. All computations were carried out using freely available code provided in the Brain Connectivity Toolbox (https://sites.google.com/site/bctnet/) [84].

First, we note a subtle but important distinction between network communication models and measures. A Network communication model (e.g., shortest path routing) delineates a strategy or algorithm to transfer information between node pairs. In turn, a network communication measure (e.g., shortest path efficiency) quantify, from a graph-theoretical standpoint, the efficiency of information transfer achieved by a given communication model. For simplicity, we used “model” throughout this paper to refer to both network communication models and measures.

We also note that certain communication measures are inherently asymmetric, in that *C_asy_*(*i, j*) ≠ *C_asy_*(*j,i*). While this asymmetry contains meaningful information on signaling properties of nervous systems [8], in the present study we consider symmetric communication matrices given by *C*(*i, j*) = (*C_asy_*(*i, j*) + *C_asy_*(*j, i*))/2. This simplification allows us to take into account only the upper triangle of *C*, substantially reducing the dimensionality of our predictive models and contributing to the computational tractability of our analyses.

#### Shortest path efficiency

Shortest path routing proposes that neural signaling takes place along optimally efficient paths that minimize the sum of connection lengths traversed between nodes. Let 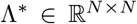 denote the matrix of shortest path lengths, where Λ^*^(*i, j*) = *L*(*i, u*) + … + *L*(*v, j*) and {*u*,…, *v*} is the sequence of nodes visited along the shortest path between nodes *i* and *j*. Shortest path efficiency was defined as SPE = 1/Λ* [9]. We computed binary (*SPE_bin_*), weighted (*SPE_wei_*), and distance (*SPE_dis_*) shortest path efficiency matrices based on the *L_bin_, L_wei_, L_dis_* connection length matrices, respectively.

#### Navigation efficiency

Navigation routing identifies communication paths by greedily propagating information based on a measure of node (dis)similarity [11]. Following previous studies on brain network communication, we used the Euclidean distance between region centroids to guide navigation [8, 12]. Navigating from node *i* to node *j* involves progressing to *i*’s neighbor that is closest in distance to *j*. This process is repeat until *j* is reached (successful navigation) or a node is revisited (failed navigation). Successful navigation path lengths are defined as Λ(*i, j*) = *L*(*i, u*) + … + *L*(*v, j*), where {*u*,…, *v*} is the sequence of nodes visited during the navigation from *i* to *j*. Failure to navigate from *i* to *j* yields Λ(*i, j*) = ∞. Navigation efficiency was defined as *NE* = 1/Λ. Binary (*NE_bin_*), weighted (*NE_wei_*), and distance (*NE_dis_*) navigation efficiency matrices were computed based on the *L_bin_, L_wei_, L_dis_*, respectively.

#### Diffusion efficiency

Diffusion efficiency models neural signaling in terms of random walks. Let 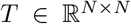 denote the transition matrix of a Markov chain process unfolding on the connection weight matrix *W*. The probability that a naive random walker at node *i* will progress to node *j* is given by 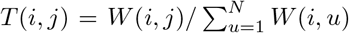. The mean first passage time *H*(*i, j*) quantifies the expected number of intermediate regions visited in a random walk from *i* to *j* (details on the mathematical derivation of *H* from *T* are given in [13, 85, 86]). Diffusion efficiency is defined as *DE* = 1/*H*, thus capturing the efficiency of neural communication under a diffusive propagation strategy [13]. Binary (*DE_bin_*), weighted (*DE_wei_*), and distance (*DE_dis_*) diffusion efficiency matrices were computed based on the *W_bin_, W_wei_, W_dis_*, respectively.

#### Search Information

Search information is derived from the probability of random walkers serendipitously traveling along the shortest paths between node pairs [14]. Let Ω(*i, j*) = {*u*,…, *v*} be the sequence of nodes along the shortest path from node *i* to node *j* computed from the connection length matrix *L*. The probability of a random walker starting at *i* reaches *j* via Ω(*i, j*) is given by *P*(Ω(*i, j*)) = *T*(*i, u*) × … × *T*(*v, j*), where *T* is the previously defined transition probability matrix computed from *W*. We defined search information as *SI*(*i, j*) = log_2_(*P*(Ω(*i, j*))) [6, 8]. This definition quantifies how accessible shortest paths are to naive random walkers, capturing the degree to which efficient routes are hidden in network topology. Note that the computation of search information depends both on *L*—for the identification of shortest paths—and *W*—for the simulation of random walk processes. We used *W_wei_* combined with *L_bin_, L_wei_*, and *L_dis_* to compute, respectively, binary, weighted, and distance versions of search information.

#### Communicability

Communicability models neural signaling as a diffusive process unfolding simultaneously along all possible walks in a network [15]. Communicability between nodes *i* and *j* is defined as the weighted sum of the total number of walks between them, with each walk weighted by its length (i.e., number of connections traversed). In the binary case, this yields 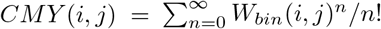. In the limit *n* → ∞, this sum converges to *CMY*(*i, j*) = *e*^*W_bin_*(*i, j*)^. Non-binary connection weight matrices are typically normalized as 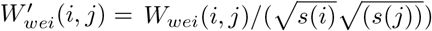 prior to the computation of communicability to attenuate the influence of high strength nodes [16], where 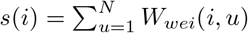 is the total strength of node *i*. We used *W_bin_*, 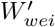 and 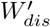 to compute, respectively, binary, weighted and distance-based versions of communicability.

### Functional connectivity data

Minimally preprocessed resting-state functional MRI data from the same 889 individuals was also obtained from the HCP. Participants were scanned twice (right-to-left and left-to-right phase encodings) on two separate days, resulting in a total of four sessions per individual. In each session, functional MRI data was acquired for a period of 14m33s with 720ms TR. Further details on resting-state functional MRI data collection and preprocessing as described in [78, 87]. Functional activity in each of *N* = 68, 360 regions was computed by averaging the signal of all vertices comprised in the region. Pairwise Pearson correlation matrices were computed from the regional time series of each session, resulting in four matrices per participant. For each participant, the four matrices were averaged to yield a final *N* × *N* FC matrix. Group-level functional connectomes were computed by averaging the FC matrices of all subjects.

### Behavioral dimensions

Information on HCP behavioral protocols and procedures is described elsewhere [88]. A total of 109 variables measuring alertness, cognition, emotion, sensory-motor function, personality, psychiatric symptoms, substance use and life function were selected from the HCP behavioral dataset [19]. Selected items consisted of raw (age- and gender-unadjusted), total or subtotal assessment scores. The set of 109 measures was submitted to an independent component analysis (ICA) pipeline in order to derive latent dimensions summarizing orthogonal dimensions of behavioral information. This procedure contributed to the computational tractability of our analyses by enabling behavioral inferences to be performed on a small set of data-driven components, rather than restricted to arbitrarily selected measures.

Behavioral dimensions were computed as follows. A rank-based inverse Gaussian transformation [89] was used to normalize continuous behavioral variables (87 of 109). Age and gender were regressed out from all behavioral items. ICA was performed on the resulting residuals using the FastICA algorithm [90] implemented in the icasso MALTLAB package [91]. Participants were sampled with replacement to generate a total of 500 bootstrap samples. ICA was independently performed on each sample with randomly selected initial conditions. Agglomerative clustering with average linkage was used to derive consensus clusters of independent components across different bootstrap samples and initial conditions. This procedure, including bootstrapping and randomization of initial conditions, was repeated for 10 trials of a set of candidate ICA models ranging from 3 to 30 independent components. The best number of components was estimated based on the reproducibility across the 10 trials by means of a cluster quality index. Clearly separated clusters indicate independent components were consistently and reliably estimated, despite being computed based on different bootstrap samples and initial conditions. This criterion identified the five component model as the most robust and parsimonious set of latent dimensions. The obtained components were visualized as word clouds with font size proportional to the contribution of each original behavioral variable (see Fig. S14 in [19]). This enabled the characterization of the five dimensions as cognitive performance, illicit substance use, tobacco use, personality and emotion traits, and mental health. Further details on the computation of the behavioral dimensions are provided in [19].

### Behavioral prediction framework

Let 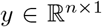 be a vector of response variables corresponding to a given behavioral dimension, where *n* = 899 is the number of individuals in our sample. Let 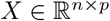 be a matrix of *p* explanatory variables corresponding to the upper triangle of vectorized communication matrices 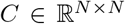, so that *p* = *N*(*N* − 1)/2. We applied two independent statistical models to predict *y* from *X*: lasso regression and a regression model based on network features identified by the NBS. These models implement different strategies of feature selection aimed at identifying a parsimonious set of variables in *X* to predict *y*.

The data was split into train and test sets to perform 10-fold cross-validation. The family structured in the HCP dataset was taken into account by ensuring that individuals of the same family were not separated between train and test sets [54]. Sensitivity to particular train-test data splits was addressed by repeating the 10-fold cross-validation 10 times. The same train and test sets were used for lasso and NBS regressions. Model parameter estimation was performed exclusively on train sets while model performance was assessed exclusively on test sets.

#### Lasso regression

Let {*X_a_, y_a_*} and {*X_e_, y_e_*} denote a split of {*X, y*} into train and test sets, respectively. We used lasso regression [22] to compute *β* as

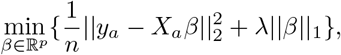

where 0 ≤ λ ≤ 1 is a feature selection parameter controlling model complexity. We systematically evaluated 100 logarithmically-spaced values of λ ranging from 0.001 to 1. Model fit was evaluated as the Pearson correlation coefficient between 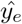 and *y_e_*, where 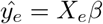. For each value of λ, model performance was computed as the average model fit over 100 pairs of train and test sets (10 repetitions of 10-fold splits). To avoid trivial cases for which maximum performance is achieved for ||*β*||_1_ = 0, we selected the smallest λ leading to model performance within one standard deviation of the maximum performance.

#### Network-based statistics regression

The NBS identifies sets of connected components that explain significant interindividual variation in a response variable [23]. We used the NBS as a feature selection technique to identify behaviorally relevant groups of connections. We then fit a regression model to the average connection weight of the selected connections in order to predict behavior. Importantly, connected components were identified exclusively in training sets, while prediction accuracy were computed based on held-out test sets.

Let {*X_a_, y_a_*} and {*X_e_, y_e_*} denote a split of {*X, y*} into train and test sets, respectively. The cross-validated predictive utility of NBS connected components was computed as follows. For each column of *X_a_* (corresponding to the value of a connection in the upper triangle of a communication matrix *C*(*i, j*) across subjects in the train set), the inter-individual Pearson correlation between *C*(*i, j*) and *y_a_* was computed. Connections for which statistical association strength exceeded a t-statistic threshold *t* > |3| were grouped into sets of positive (*t* > 3) and negative (*t* < −3) connected components. This procedure was repeated for 1000 random permutations of *y_a_*, and the likelihood of observing positive and negative connected components as large as empirical ones was assessed using a non-parametric test. Further details on the NBS are found in [23].

Let Γ^+^ and Γ^−^ be, respectively, the largest positive and negative connected components identified by the NBS based on the train data {*X_a_,y_a_*}. We computed 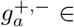 as the average weight of connections belonging to the connected component Γ^+,−^:

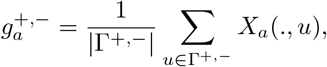

where |Γ^+,−^| indicates the number of connections comprising the connected component. We defined the matrix 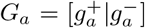. Therefore, 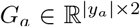 contains the average weight of connections identified as positively and negatively associated with the behavioral dimension *y* for subjects in the train set {*X_a_,y_a_*}. Using a bivariate linear regression model, we computed the coefficients *β* such that

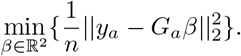

Analogously, we computed the average weight of connected components in the test set

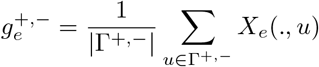

and 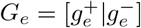. Finally, behavioral predictions were computed as 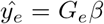 and out-of-sample prediction accuracy was evaluated as the Pearson correlation coefficient between 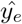 and *y_e_*. This procedure was repeat for 100 pairs of train and test sets (10 repetitions of 10-fold splits). In cases where no significant component was identified by the NBS (|Γ^+,−^| = 0), model performance was set to 0.

### Functional connectivity prediction framework

FC prediction utility was determined via Spearman correlation between empirical FC and analytically derived communication matrices. Therefore, none of the models and measures used to infer FC required training, statistical estimation of weights or parameter tuning (an advantage over other classes of high-order models). Hence, we oftentimes adopted the term FC “prediction” even though predictive utility was not assessed out-of-sample [6].

### Overall predictive utility rankings

Overall ranking of communication model predictive utility were computed by averaging rankings obtained for behavioral and functional analyses, which were given equal weight on the averaging process. The overall ranking for thresholded *N* = 360 connectomes was computed as the weighted average of rankings for (i) lasso behavioral predictions (0.25 weight), (ii) NBS behavioral predictions (0.25 weight), and (iii) FC predictions (0.5 weight). Rankings for other connectome mapping pipelines were computed by averaging, with equal weight, lasso behavioral predictions and FC predictions.

## SUPPLEMENTARY INFORMATION

### Note 1: Behavioral predictive utility for the cognition and tobacco use dimensions

We have characterized the predictive utility of connectivity measures and communication models by considering a pooled prediction accuracy between the cognition and tobacco use dimensions. As we have seen, these behavioral dimensions led to the most accurate and consistent predictions in our sample. Considering the average prediction accuracy across the two traits facilitated the comparison of communication models by providing us with a single score on which to evaluate prediction accuracy. However, this approach may potentially obscure nuanced phenotype-specific relationships between brain and behavior. Indeed, we observed no correlation between the prediction accuracies obtained for cognition and tobacco use (Spearman rank correlation *p* = 0.49,0.69 for lasso and NBS, respectively). Therefore, in this section, we sought to separately examine behavioral predictions for the cognition and tobacco use dimensions.

We observed that the outstanding performance of FC was mostly due to the cognition dimension (Fig. S4 and S5; this can also be observed in Fig. 2c,d). This is in line with several findings on the relationship between cognitive processes and the architecture of functional networks [46]. Interestingly, while FC still yielded top-ranking predictions of tobacco use, several communication models showed comparable predictive utility in this dimension (Fig. S6 and S7; again evident in Fig. 2c,d). For instance, navigation was the best predictor of tobacco use, with binary and distance navigation occupying the first positions under lasso and NBS, respectively. Although none of the communication models statistically outperform FC, this points towards a tighter margin of difference between the utility of structural and functional measures in predicting behavioral phenotypes not directly related to cognition.

Another interesting observation was that weighted search information was the best communication model in predicting cognition, but showed near bottom-ranking predictive utility of tobacco use, which resulted in a moderate performance when combining predictions from both behavioral dimensions. Along similar lines, we observed that rankings of connection weight definitions were sensitive different behavioral dimensions and prediction methods, painting an unclear picture of what weighting schemes best contribute to predict human behavior from structural connectomes.

Therefore, collectively, our results did not point towards a single communication model as the best predictor of human behavior. Despite the overall good performance of communicability and navigation, our observations indicate that different communication models may be better suited to predict different behavioral dimensions, possibly suggesting the presence of behavior-specific signal-ing mechanisms in the human brain. Importantly, across the multiple analyses performed, our results consistently suggested that network communication models, in particular communicability and navigation, improve the behavioral predictive utility of the human connectome.

### Note 2: Additional analyses of structure-function coupling

We further examined structure-function relationships by stratifying FC predictions across anatomically connected and unconnected regions pairs (Fig. S9a). In accordance to previous work [6, 12], associations to FC were stronger for connected regions. Despite these changes in association strength, rankings of communication models in terms FC predictions were consistent across the scenarios explored (Spearman rank correlation *r* = 0.90, 0.68, 0.85 and *p* = 8 × 10-7, 3 × 10-3,0 between connected and all, all and unconnected, and connected and unconnected region pairs, respectively). Interestingly, certain communication models outperformed SC for connected regions, suggesting that indirected polysynaptic signaling maybe relevant for communication even in the presence of direct anatomical links.

Estimates of structure-function coupling depend on accurate reconstruction of structural connectomes. However, white matter tractography is prone to a number of known biases, of which the underestimation of inter-hemispheric connections is an important concern [73]. To attenuate this issue, past studies have focused on intra-hemispheric characterization of structure-function coupling [41, 56]. Focusing on the right hemisphere, we found that, for all communication models, FC associations were stronger compared to whole-brain estimates (Fig. S9b). Importantly, the functional predictive utility ranking of communication models remained consistent (Spearman rank correlation *r* = 0.93 and *p* = 0 between whole-brain and right hemisphere rankings). This suggests that our analyses may provide a meaningful ranking of signaling strategies despite shortcomings in connectome mapping techniques.

**FIG. S1.**
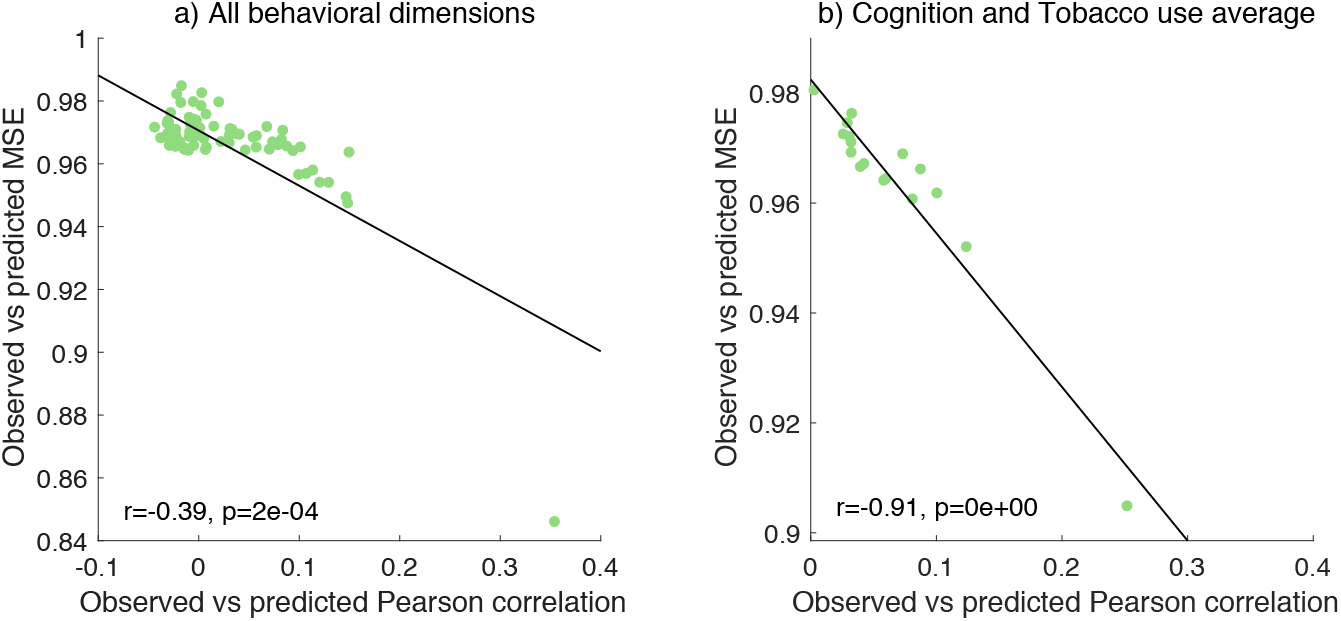
Comparison between behavioral prediction accuracies computed using Pearson correlation and mean square error evaluated using Spearman rank correlation (Lasso regression, *N* = 360 thresholded connectomes). The least squares line is shown in black.

**FIG. S2.**
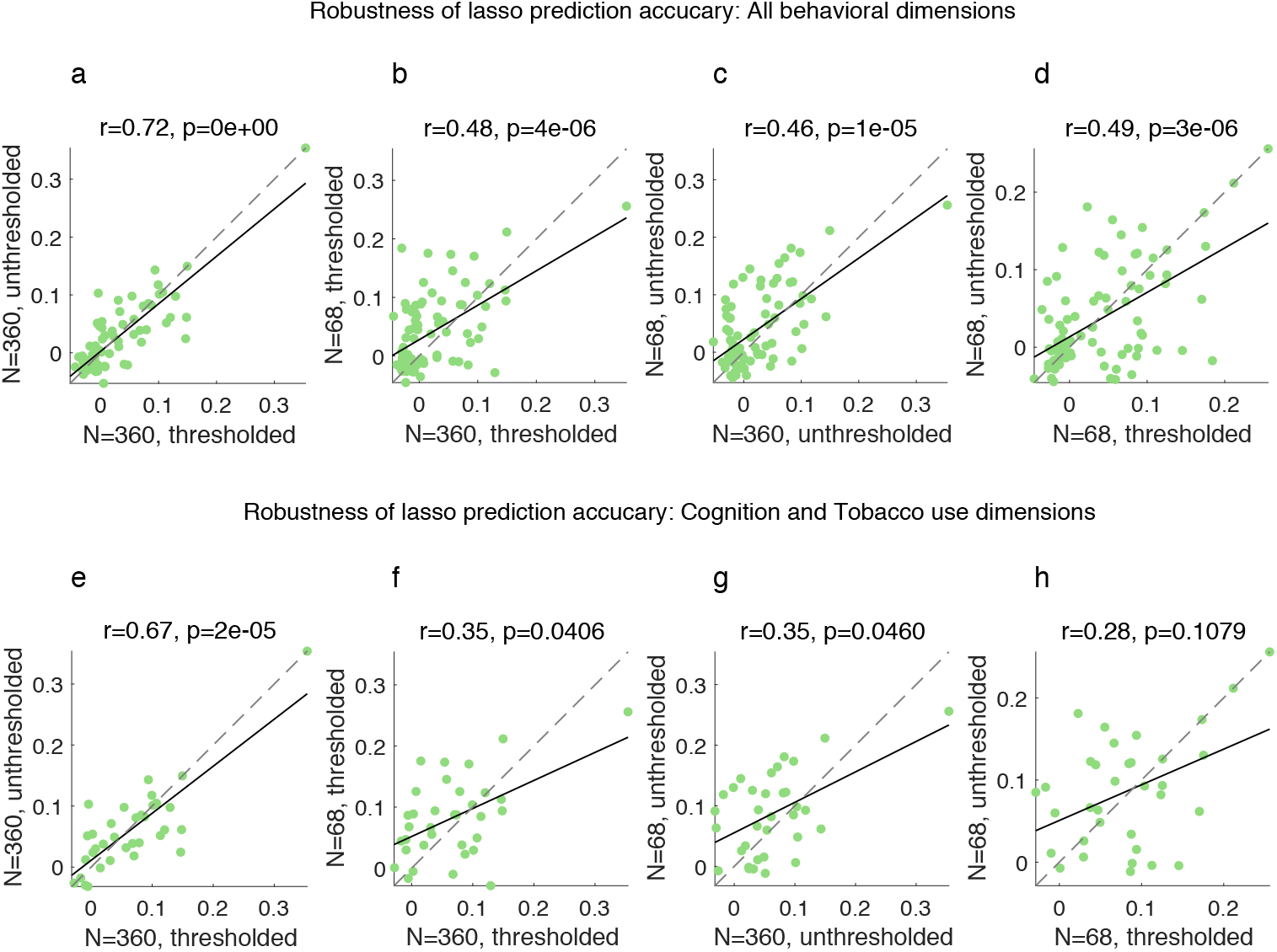
Comparison between behavioral prediction accuracies across multiple connectome mapping pipelines (Lasso regression). Scatter plots show the least squares and *x* = *y* lines are shown in solid black and gray dashed traces, as well as the Spearman rank correlation coefficient and *p*-value.

**FIG. S3.**
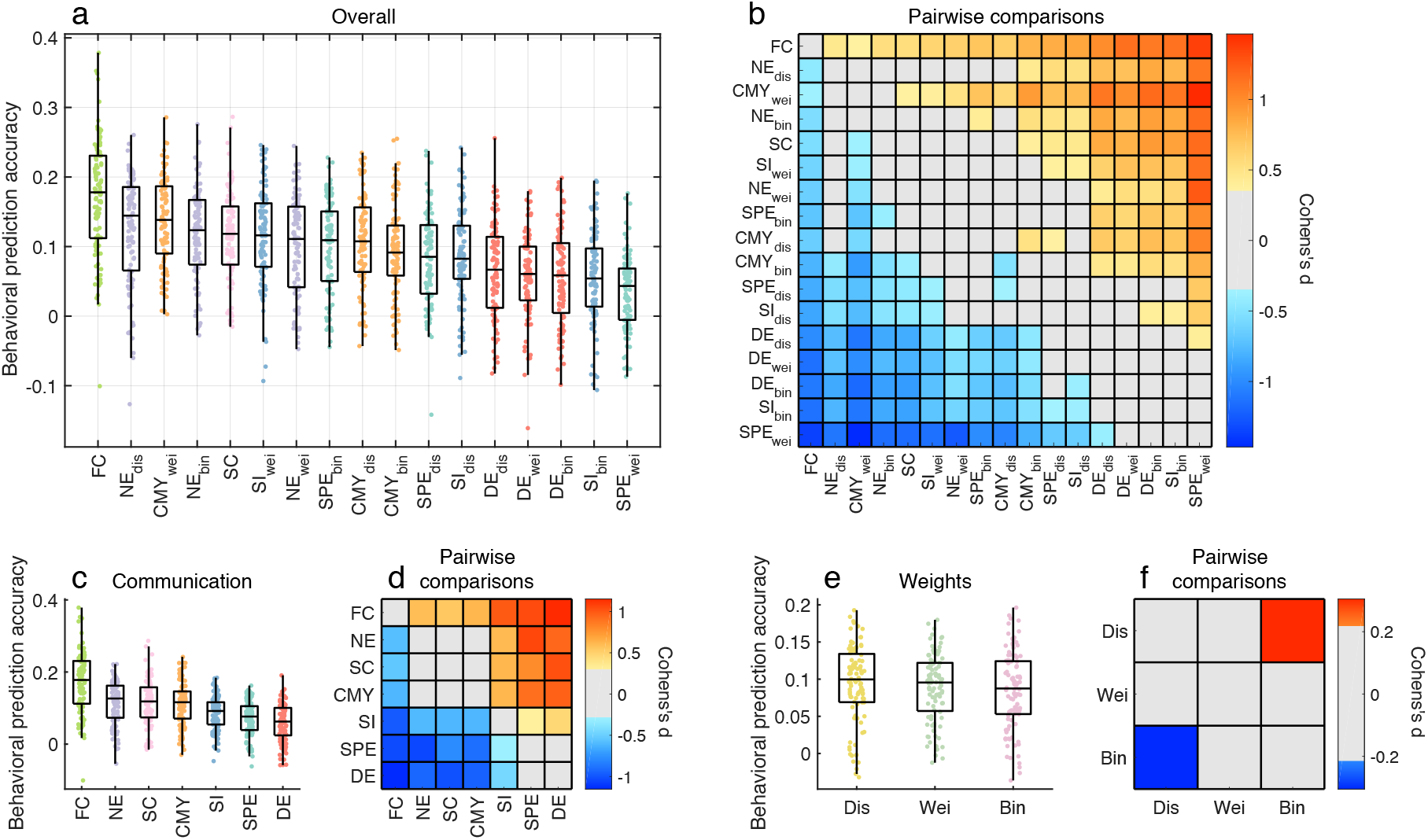
Comparison of the behavioral predictive utility of connectome communication models (NBS, *N* = 360 thresholded connectomes, average cognition and tobacco use prediction accuracies). It is worth noting that the NBS feature selection process is better suited to sparse graphs [23], which could confer an advantage to sparse SC matrices over fully-connected communication and FC matrices.

**FIG. S4.**
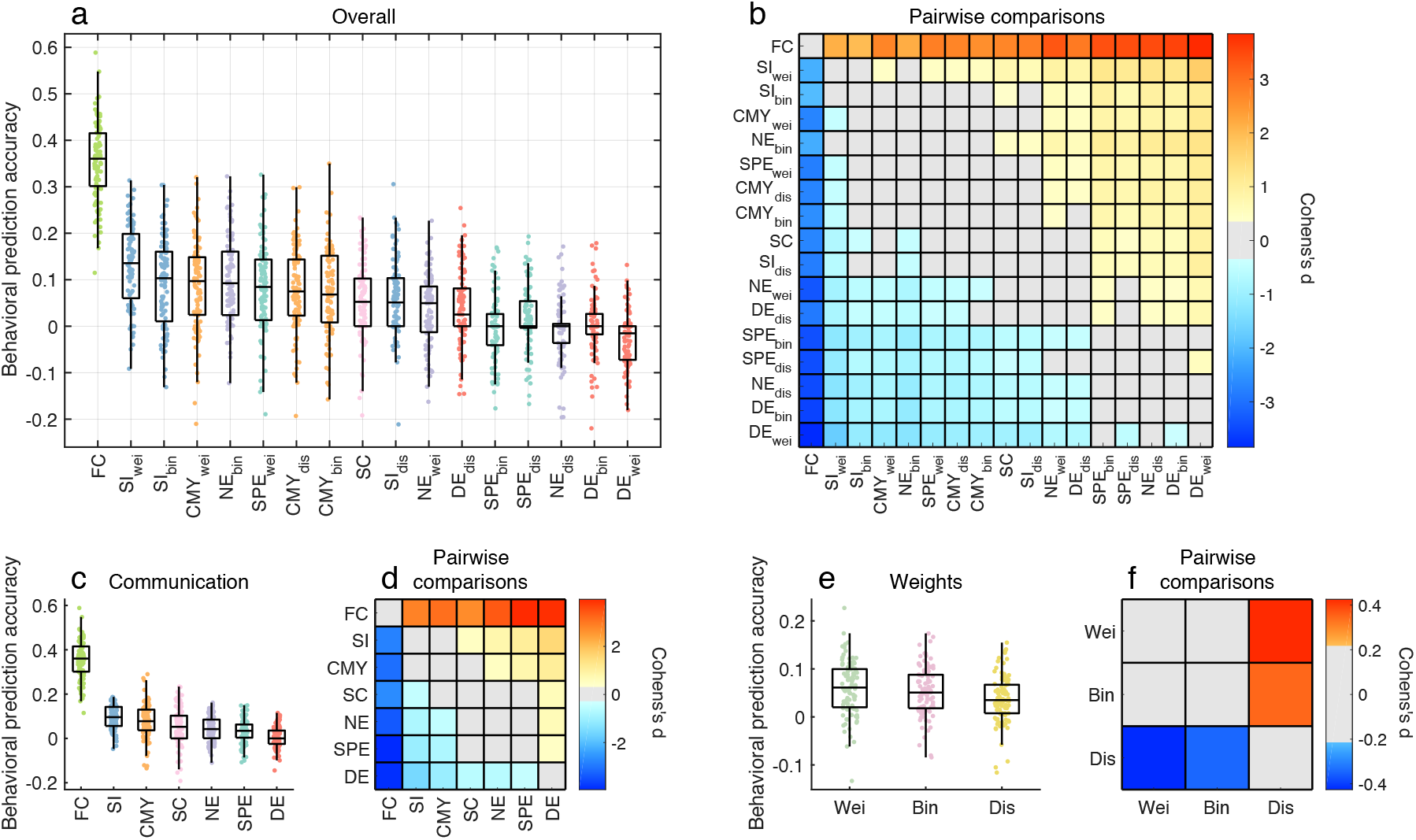
Comparison of the behavioral predictive utility of connectome communication models (Lasso regression, *N* = 360 thresholded connectomes, cognition prediction accuracy).

**FIG. S5.**
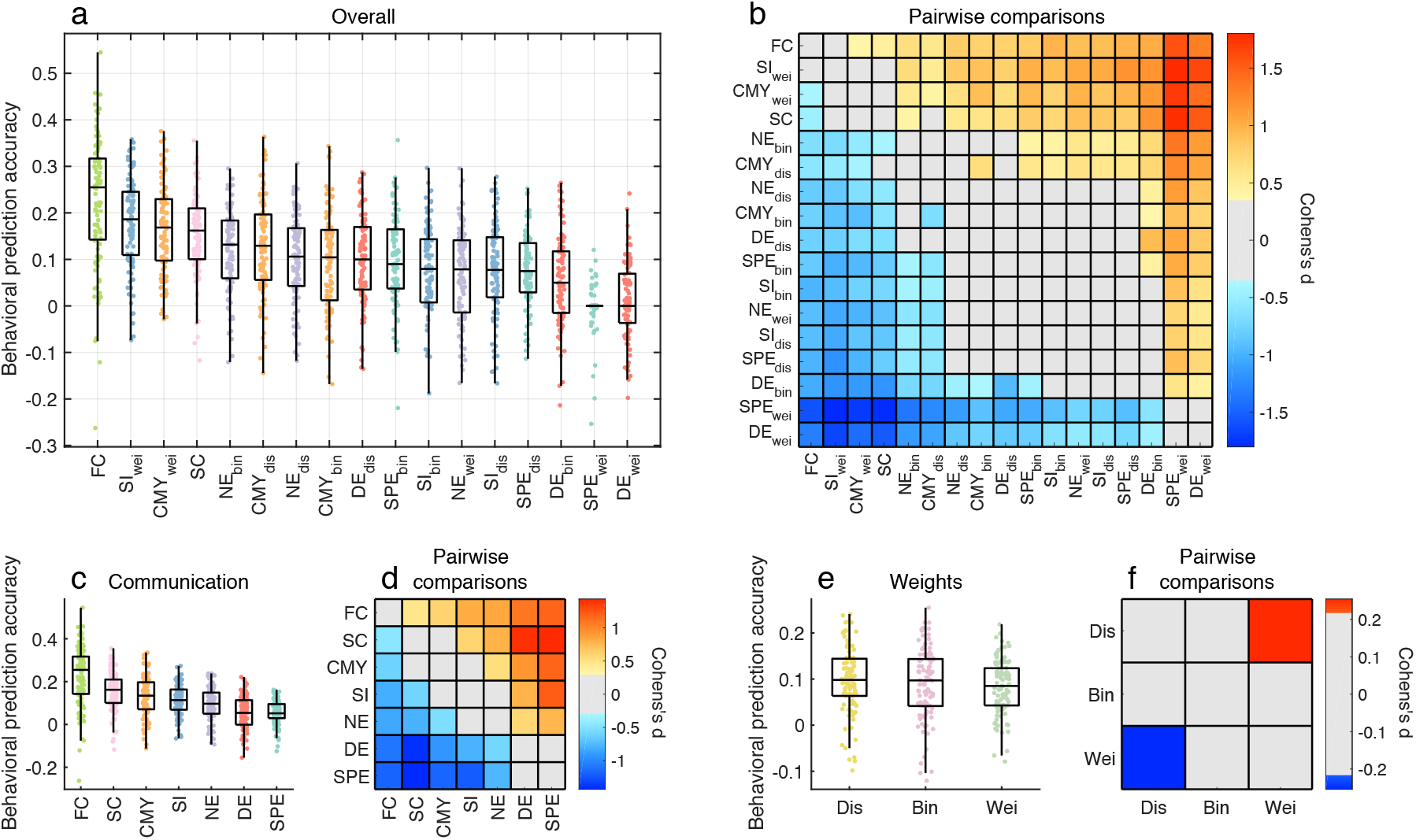
Comparison of the behavioral predictive utility of connectome communication models (NBS, *N* = 360 thresholded connectomes, cognition prediction accuracy)

**FIG. S6.**
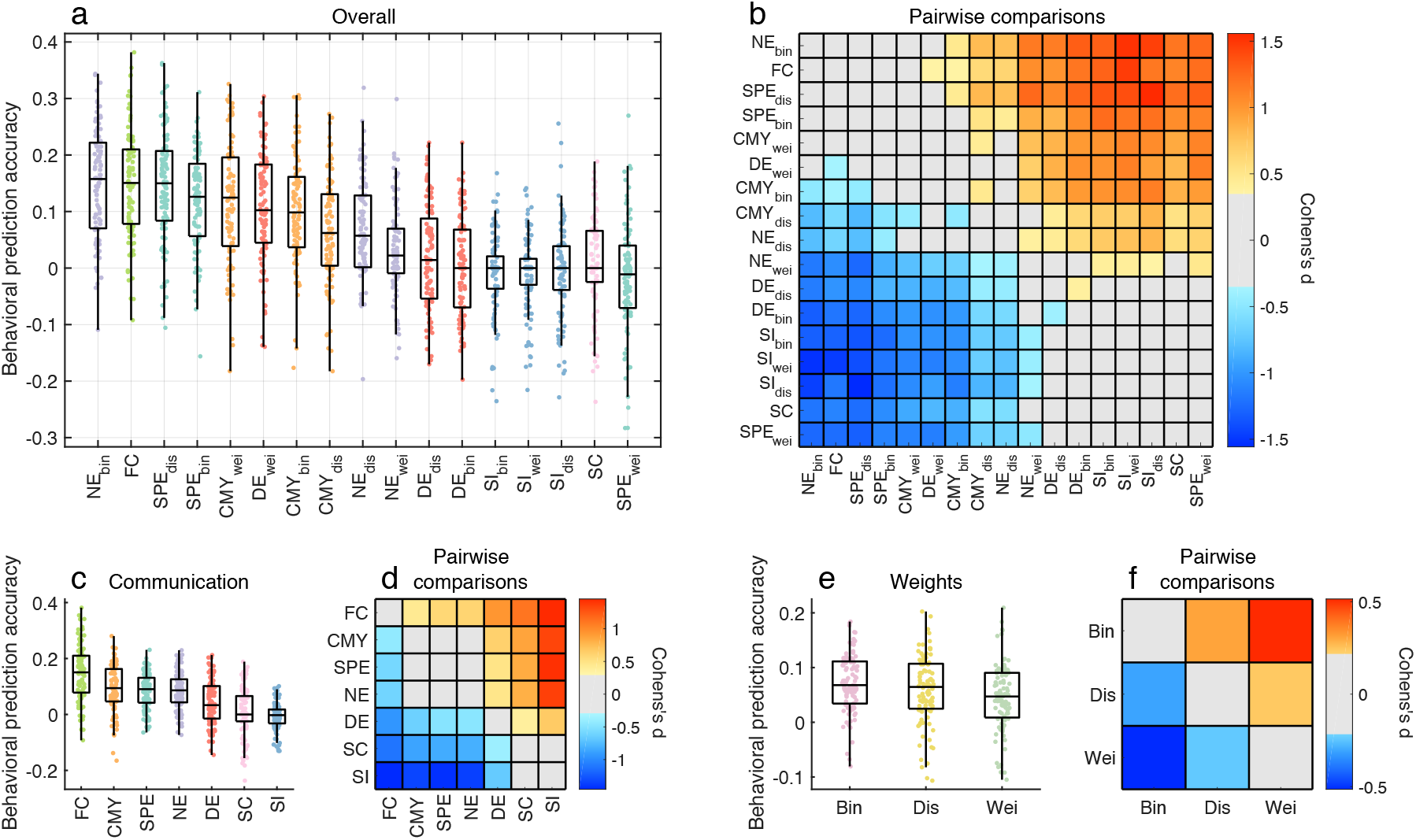
Comparison of the behavioral predictive utility of connectome communication models (Lasso regression, *N* = 360 thresholded connectomes, tobacco use prediction accuracy)

**FIG. S7.**
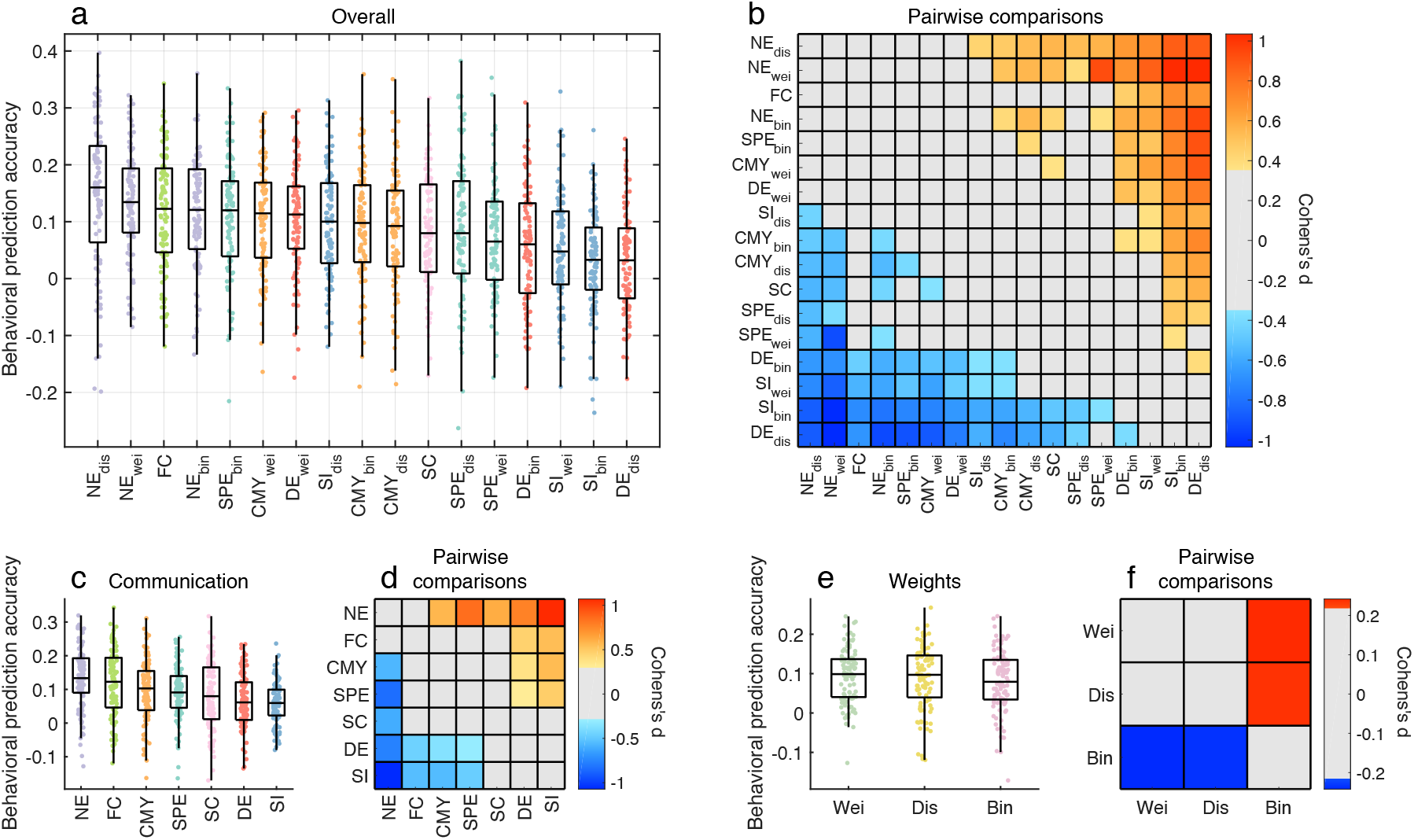
Comparison of the behavioral predictive utility of connectome communication models (NBS, *N* = 360 thresholded connectomes, tobacco use prediction accuracy

**FIG. S8.**
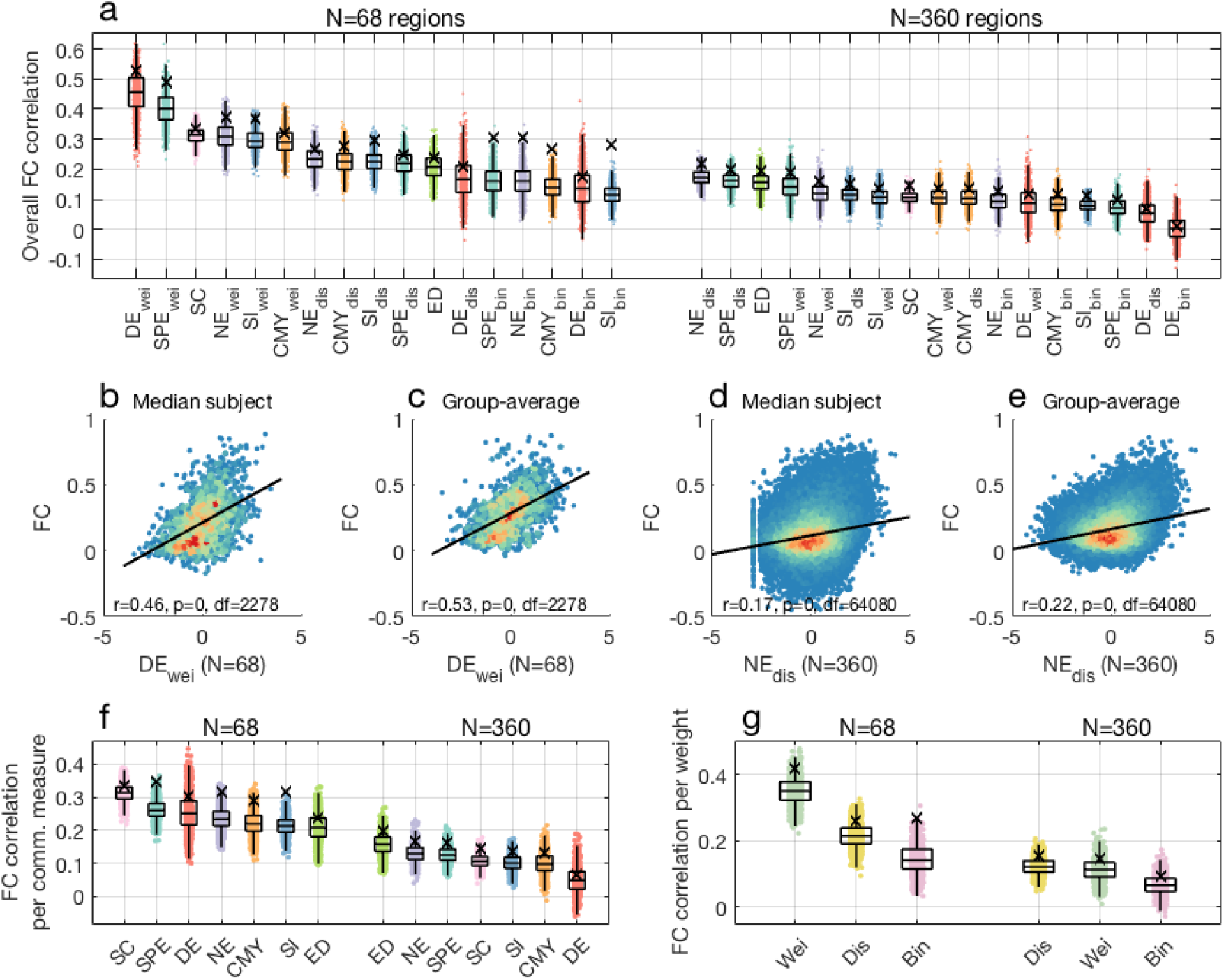
Structure-function coupling across connectome communication models (*N* = 68, 360 unthresholded connectomes).

**FIG. S9.**
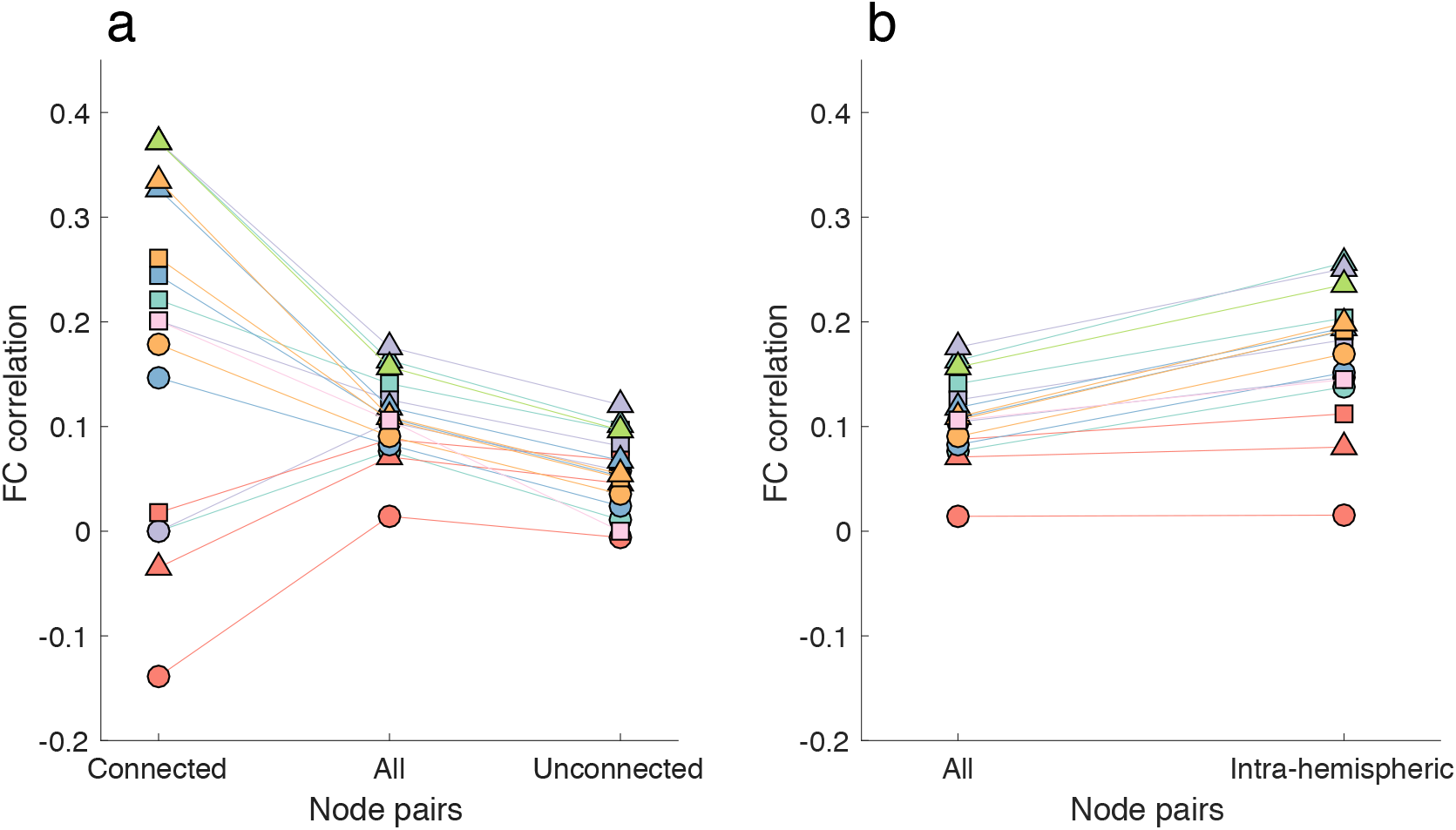
Stratification of structure-function coupling across connectome communication models (*N* = 360 thresholded connectomes). Markers indicate median (across subjects) Spearman correlation between FC and communication models. Squared, circular and triangular markers denote models computed on weighted, binary and distance connectomes, respectively. Marker colors denote different FC predictors. Teal: shortest paths, violet: navigation, red: diffusion, blue: search information, orange: communicability, pink: SC, and green: Euclidean distance. **(a)** Structure-function coupling anatomically connected (left), all (center) and anatomically unconnected (right) region pairs. **(b)** Structure-function coupling for whole-brain (left) and right-hemisphere (right) connectomes.

**FIG. S10.**
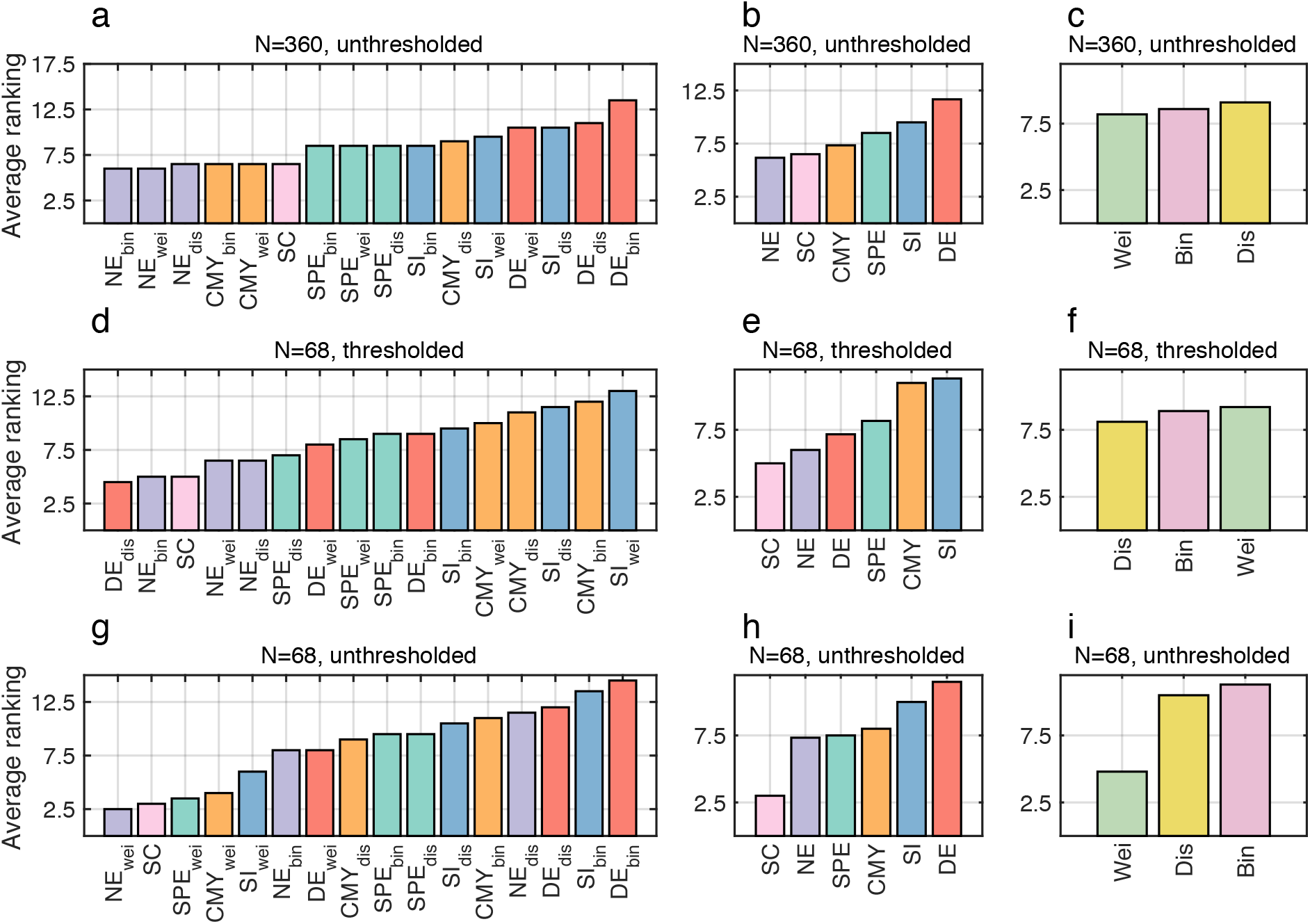
Overall ranking of communication models (multiple connectome mapping pipelines). Communication models and SC were ranked according to averaged behavioral and functional predictions. **(a,d,g)** Overall predictive utility ranking. Rankings shown in (a,d,g) grouped by **(b,e,h)** communication models and **(c,f,i)** connection weight definitions.

